# iCLIP3: A streamlined, non-radioactive protocol for mapping protein-RNA interactions in cellular transcripts at single-nucleotide resolution

**DOI:** 10.64898/2026.03.01.708747

**Authors:** Vladimir Despic, Melina Klostermann, Anna Orekhova, Mikhail Mesitov, Anke Busch, Kathi Zarnack, Julian König, Michaela Müller-McNicoll

## Abstract

UV-C crosslinking and immunoprecipitation (CLIP)–based methods are the gold standard for identifying direct RNA binding protein (RBP) interaction sites on cellular RNA *in vivo*. Here, we describe individual-nucleotide resolution CLIP version 3 (iCLIP3), an optimized protocol for generating transcriptome-wide maps of RBP–RNA interaction sites at single-nucleotide resolution from low-input material. iCLIP3 introduces several key improvements over previous iCLIP variants, including rapid and safe infrared-based visualization of RBP–RNA complexes, silica column-based RNA isolation, and the incorporation of TruSeq adapter sequences with unique dual indexing. These modifications streamline library preparation, facilitate multiplexing, and enable concurrent sequencing of iCLIP3 libraries alongside unrelated RNA-seq libraries. In addition, we provide a detailed bioinformatics workflow for identifying crosslinking events and defining RBP binding sites. The complete protocol can be performed within 4–5 days, offering a robust and efficient approach for high-resolution mapping of RBP–RNA interactions.

**Highlights:**

- 3′ RNA labeling with pCp-IR750 enables rapid infrared visualization of RBP–RNA complexes
- Silica column-based RNA isolation improves efficiency and reproducibility
- TruSeq adapters with unique dual indexing enable flexible and scalable multiplexing
- iCLIP3 generates high-quality libraries from ultra-low amounts of cellular material

## BEFORE YOU BEGIN

### Overview

UV-C crosslinking and immunoprecipitation (CLIP)-based techniques provide a powerful approach for transcriptome-wide mapping of RNA binding protein (RBP)–RNA interactions at nucleotide resolution in living cells. These methods rely on the ultraviolet (UV)-induced covalent crosslinking of RBPs to their target RNAs—thereby preserving direct protein–RNA interactions *in vivo*—followed by immunoprecipitation (IP) of the crosslinked ribonucleoprotein complexes and identification of associated RNA fragments by high-throughput sequencing (Ule et al., 2003; Lee and Ule, 2018).

Several CLIP variants have been developed to enhance crosslinking efficiency and library complexity or provide complementary information, including high-throughput sequencing CLIP (HITS-CLIP) (Licatalosi et al., 2008), photoactivatable ribonucleoside-enhanced CLIP (PAR-CLIP) (Hafner et al., 2010), and enhanced CLIP (eCLIP) (Van Nostrand et al., 2016; Blue et al., 2022). A defining feature of individual-nucleotide resolution CLIP (iCLIP) is its ability to capture the precise positions of protein–RNA contacts by exploiting reverse transcriptase truncation sites to achieve nucleotide resolution (König et al., 2010; Huppertz et al., 2014; Buchbender et al., 2020). We have used iCLIP to pinpoint differences in the binding preferences of RBPs in different cellular compartments (Brugiolo et al., 2017), upon cellular stress (de Oliveira Freitas Machado et al., 2023), between closely related family members (Müller-McNicoll et al., 2016), or between disease-relevant mutants (Zimmann et al., 2025) as well as to dissect complex assembly at 3′ splice sites (Zarnack et al., 2013; Sutandy et al., 2018; Ebersberger et al., 2023) and protein binding to RNA modifications (Zhou et al., 2024). Collectively, these approaches provide complementary strategies for the systematic and comprehensive, transcriptome-wide characterization of RBP–RNA interactions at single-nucleotide resolution.

In iCLIP, immunopurified RBP–RNA complexes are denatured, separated by SDS–PAGE, and transferred to a nitrocellulose membrane. RNA fragments are isolated, and the crosslinked protein is removed via protease digestion but leaves a short peptide at the crosslink site. During reverse transcription, this residual peptide induces premature termination of cDNA synthesis at the crosslink site. Sequencing of these truncated cDNAs enables the precise, nucleotide-level mapping of RBP binding sites across the transcriptome.

#### Innovation

iCLIP3 builds upon the previously published iCLIP2 protocol (Buchbender et al., 2020) and introduces several methodological improvements that simplify the workflow, enhance safety, and increase compatibility with standard sequencing pipelines.

First, iCLIP3 replaces radioactive 5′-end RNA labeling with infrared 3′-end RNA labeling using the pCp-IR750 dye, enabling non-radioactive visualization of RBP–RNA complexes (**Fig. 1**). This labeling approach allows direct assessment of RNase I–mediated RNA fragmentation and assessment of RNA fragment size distributions within immunopurified complexes prior to library preparation.

**Figure 1.**
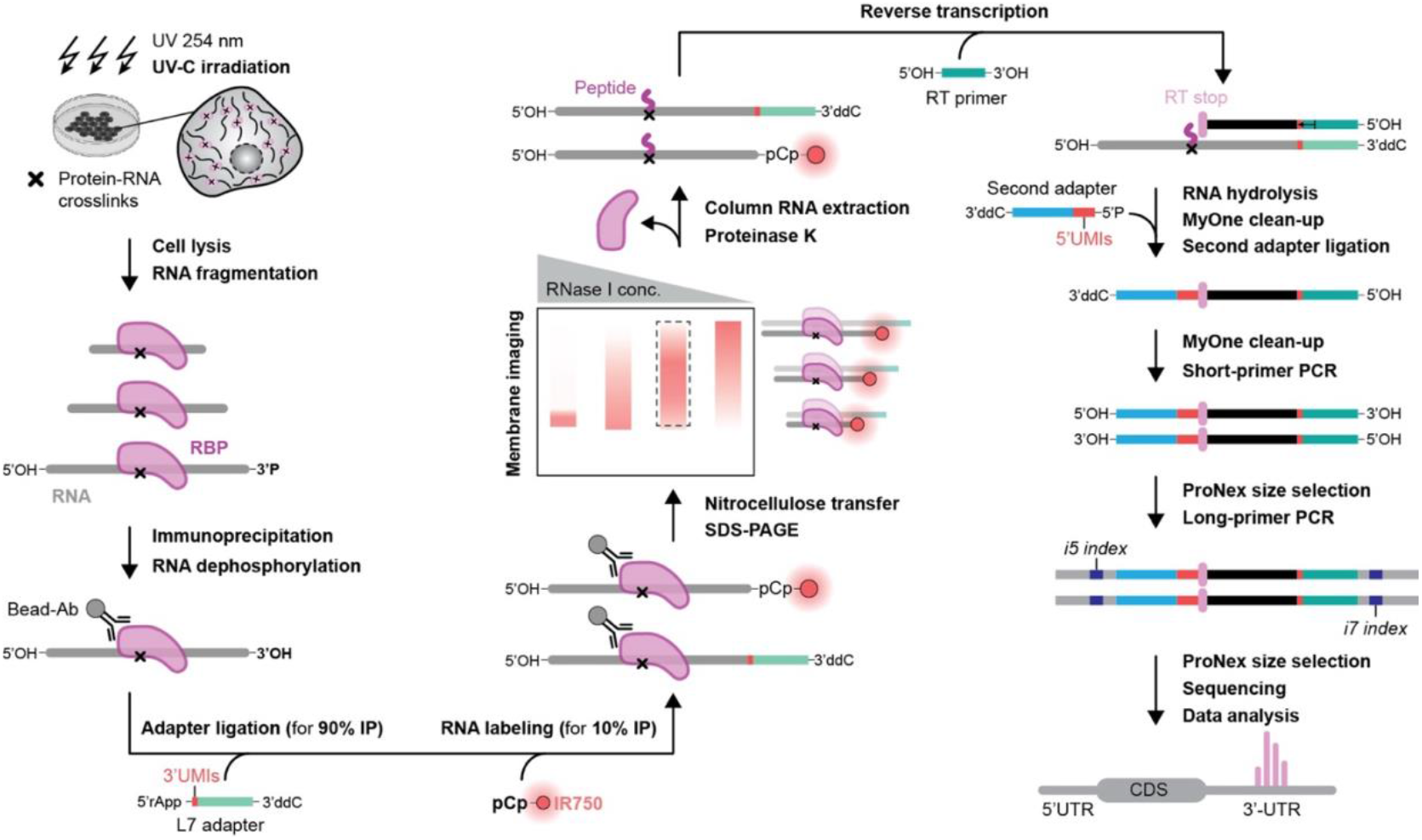
Overview of the iCLIP3 library preparation workflow. Living cells are irradiated with 254-nm UV light to induce covalent crosslinking between RNA and RBPs, followed by cell lysis. RNA is partially fragmented with RNase I in cleared lysates of defined protein amount and concentration. The RBP of interest is then purified using a specific antibody coupled to magnetic Protein A or G beads. 3′-ends of the immunopurified RNA fragments are dephosphorylated on the beads using T4 polynucleotide kinase (PNK) enzyme. L7 linker, a 3′-end adapter, is ligated on beads to the RNA component of the 90% of purified protein–RNA complexes, whereas pCp-IR750 is independently ligated to the RNA component of the remaining 10%. L7 linker contains unique molecular identifiers (UMIs) in a form of three random nucleotides, hereby named 3′UMIs. 3′UMIs are used to reduce sequence biases during L7 linker ligation. After extensive and stringent washes, two reactions are combined and RBP–RNA complexes are eluted from the beads. RBP–RNA complexes are resolved based on their size on the SDS-PAGE and transferred onto the nitrocellulose membrane. Complexes containing pCp-IR750 are visualized on the membrane in the Cy7 or IRDye 800 CW channels. The membrane area corresponding to the RBP–RNA complexes is cut out and RNA released from the membrane using Proteinase K. RNA is further purified using silica columns. Note that the RNA contains a short peptide, a remnant of the UV-crosslinked RBP. The RNA is reverse transcribed into cDNA, during which reverse transcriptase stalls at the peptide–RNA crosslink site, generating truncated cDNAs. The terminal nucleotide at the 3′ end of the cDNA thus marks the RBP crosslink position. RNA is subsequently hydrolyzed, and cDNA is extracted using MyOne Silane beads. A second DNA adapter is then ligated to the cDNA 3′ ends; this adapter contains nine random nucleotides, hereby named 5′UMIs. cDNA molecules containing both adapters are purified using MyOne Silane beads and subjected to pre-amplification using short P5 and P7 primers. The resulting double-stranded DNA is size-selected with ProNex beads to remove adapter dimers. Finally, pre-amplified cDNA is further amplified using long P5 and P7 primers containing dual *i5* and *i7* indices for sample multiplexing. Final iCLIP3 libraries are purified with ProNex beads to remove unused primers, pooled in equimolar ratios, and sequenced on Illumina platforms.

Second, RNA purification in iCLIP3 is performed using silica column-based isolation instead of phenol–chloroform extraction, simplifying sample handling and eliminating the use of organic solvents (**Fig. 1**). These modifications improve reproducibility and allow researchers without the access to radioactive infrastructure to perform iCLIP.

Third, iCLIP3 incorporates TruSeq-compatible Illumina adapter sequences and unique dual indexing, which permit multiplexing and simultaneous sequencing of iCLIP3 libraries together with standard RNA-seq libraries within the same sequencing run (**Fig. 1–2**).

**Figure 2.**
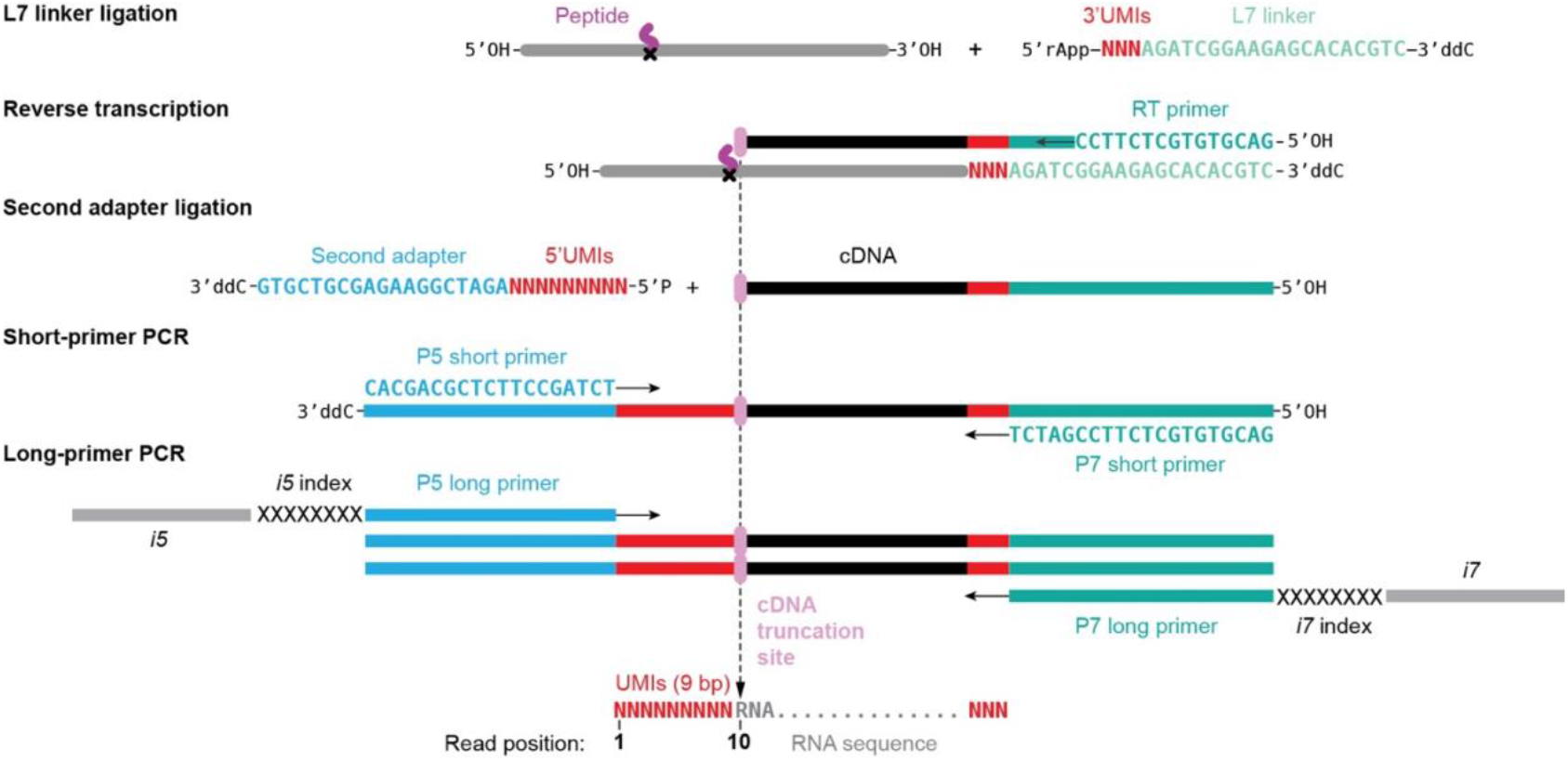
iCLIP3 oligonucleotide design. L7 linker ligation: The L7 linker is a 3′ DNA adapter with a pre-adenylated 5′ end (5′-rApp), which enables efficient ligation to the 3′ end of immunopurified RNA in the absence of ATP. A dideoxycitidine modification at the 3′ end (3′-ddC) prevents self-ligation of the linker. The L7 linker also contains three random nucleotides (NNN) at its 5′ end to reduce ligation biases. These random nucleotides are referred to as 3′ unique molecular identifiers (3′UMIs) that are appended to the 3′ end of the RNA. **Reverse transcription:** Reverse transcription (RT) is performed using an RT primer complementary to the 3′ end of the L7 linker, generating cDNA from the RNA template. **Second adapter ligation:** The second DNA adapter is phosphorylated at its 5′ end to enable ligation to the 3′ ends of truncated cDNA molecules. A 3′-ddC modification prevents adapter self-ligation. The adapter contains nine random nucleotides (NNNNNNNNN) at its 5′ end (5′UMIs) which are used for deduplication of sequencing reads. **Short-primer PCR:** Short P5 and P7 primers (P5_short and P7_short) are used for cDNA pre-amplification. **Long-primer PCR:** Long P5 and P7 primers (P5_long and P7_long) are used for the final amplification of iCLIP3 libraries. These primers contain *i5* and *i7* sequences required for cluster generation on Illumina flow cells, as well as dual 8-nucleotide indices (XXXXXXXX) for sample multiplexing.

Finally, we provide a dedicated bioinformatics workflow for iCLIP3 data analysis. This workflow uses a modified version of the pipeline racoon_clip (Klostermann and Zarnack, 2024) to identify crosslinking events (Busch et al., 2020) and the R/Bioconductor package BindingSiteFinder (Bru ggemann et al., 2021) to define RBP binding sites across biological replicates.

#### Optimizations

CLIP-based protocols require specific optimization, as their performance depends on factors such as expression levels of the RBP of interest, availability and quality of IP-grade antibodies, and the ability of the protein to directly bind RNA *in vivo*. To maximize the likelihood of successful iCLIP3 library preparation, we recommend performing the following preliminary assessments prior to initiating the full iCLIP3 workflow.

##### 1. Immunoprecipitation of the RBP of interest in iCLIP3 buffers

The suitability of the selected antibody should be evaluated by testing its ability to efficiently immunoprecipitate the protein of interest using iCLIP3 lysis and wash buffers.

##### 2. Assessment of RNA binding and optimization of RNase I-based RNA fragmentation

The number of annotated RBPs has expanded to over 1,500; however, many lack canonical RNA-binding domains and have not been validated as direct RNA binders (Hentze et al., 2025). For all proteins of interest, and particularly for non-canonical or unvalidated RBPs, we recommend the following assessments:

##### a. Verification of RNA binding *in vivo*

Confirm that the protein of interest directly associates with RNA under the chosen experimental conditions.

##### b. Optimization of RNase I-mediated RNA fragmentation

Determine the appropriate RNase I dilution required to generate RNA fragments within the optimal size range for iCLIP3 library preparation.

In addition to the main iCLIP3 library preparation protocol and the bioinformatics workflow for iCLIP3 data analysis, we provide a detailed step-by-step procedure for assessing RNA binding capacity of a protein of interest and for optimizing RNase I-based RNA fragmentation.

## PROTOCOL 1 RNA BINDING ASSAY AND OPTIMIZATION OF RNA FRAGMENTATION

### STEP-BY-STEP METHOD DETAILS

#### Cell culture and UV irradiation

##### Timing: variable, 2 days for the sample preparation

1. Cell seeding for the experiment.
  a. Grow HeLa cells in standard 1X DMEM medium supplemented with 10% FBS and 1X Penicillin-Streptomycin (Pen-Strep).
  b. One day before the experiment, seed HeLa cells onto two 100-mm dishes to achieve ∼85% confluency the following day.
2. PBS cell wash.
  a. Aspirate the culture medium from each dish.
  b. Gently wash the cells with 6 mL ice-cold PBS.
  c. Remove the PBS and add 6 mL fresh, ice-cold PBS to each dish.
3. UV irradiation.
  a. Place one cell dish containing PBS on an ice tray covered with a thin layer of water.
  b. Remove the lid and irradiate the cells once with 254-nm UV light at 150 mJ/cm^2^.
  c. Keep the second dish on ice without UV exposure as a non-irradiated control.
4. Cell harvesting.
  a. Gently scrape the cells in PBS using cell lifters.
  b. Transfer the cell suspensions into 15 mL Falcon tubes labeled UV+ (UV-irradiated) and UV– (non-UV-irradiated).
  c. Pellet the cells by centrifugation at 300 x *g* for 5 min at 4ºC. **Note**: If available, use a swing-bucket rotor to ensure pellets form at the bottom of the Falcon tubes.
  d. Carefully aspirate the PBS without disturbing the cell pellets.
  e. Snap-freeze the cell pellets on dry ice or in liquid nitrogen and store at -80ºC.

**Pause point**: Frozen cell pellets can be kept at -80ºC long term.

#### Antibody-bead coupling

**Day 1 Timing: 1–2 h**

5. Couple the antibody to magnetic Protein A or Protein G Dynabeads. **Note**: Verify the antibody’s binding preference for Protein A or Protein G Dynabeads according to the manufacturer’s recommendations.
  a. Aliquot 30 µL Protein A or G Dynabeads stock per sample into a 1.5 mL tube. **Note**: For six samples, aliquot 189 µL beads (6.3x surplus required for this protocol).
  b. Wash the beads twice with 750 µL Lysis buffer.
  c. Resuspend the beads in 300 µL Lysis buffer and add 5 µg antibody per sample specific for the RBP of interest. **Note:** For six samples, use 31.5 µg antibody in a total volume of 500 µL Lysis buffer. Lower amounts of antibody per sample may be used. All washes in the protocol are performed by pipetting unless otherwise stated.
  d. Incubate the beads on a rotating wheel at room temperature for ≥1 h (until the lysates are ready for immunoprecipitation).

#### Cell lysis and protein quantification

**Day 1 Timing: 1 h**

6. Cell lysis.
  a. Thaw frozen cell pellets on ice.
  b. Resuspend each cell pellet in 750 µL Lysis buffer supplemented with 1X Protease Inhibitors (1X PIs).
  c. Lyse the cells on ice for 10 min.
  d. **Optional:** Sonicate the lysate on ice at 10% amplitude using 5 cycles of 5 s pulses with 10 s pauses between pulses. **Note**: Sonication is recommended for nuclear proteins. If sonication is omitted, extend the lysis time to 20 min on ice.
  e. Transfer the lysates to 1.5 mL tubes.
  f. Clarify the lysates by centrifugation at 16,000 x *g* for 10 min at 4ºC.
  g. Transfer the supernatants to fresh 1.5 mL tubes.
7. Determine the protein concentration in the cleared lysates using a BCA protein quantification kit according to the manufacturer’s instructions.
8. Prepare the lysates for RNA fragmentation.
  a. Dilute the UV– and UV+ lysates to a final protein concentration of 0.5 mg/mL using Lysis buffer supplemented with 1X PIs. Prepare at least 1.6 mL of UV– and UV+ lysates at 0.5 mg/mL to proceed.
  b. Label three 1.5 mL tubes per condition as H (High), M (Medium), and L (Low).
  c. Transfer 500 µL diluted lysate (250 µg protein) into each tube and keep the lysates on ice.

#### RNA fragmentation

**Day 1 Timing: 20 min**

The iCLIP3 technology relies on short-read sequencing. In this step, RNA in the lysate is partially digested with an appropriate concentration of RNase I to generate RNA fragments in the 50–300 nucleotides (nt) size range to allow for library preparation and sequencing. Higher RNase I concentrations generating RNA fragments in the 4–20 nt size range are used for determining the efficiency and purity of the IP. The optimal RNase I concentration must be empirically determined by following the optimization guidelines below.

9. DNase treatment.
  a. Add 2 µL **TURBO DNase** to each diluted lysate.
  b. Gently invert the tubes several times to mix.
  c. Briefly spin down the samples and return them to ice.
10. RNase I digestion.
  a. Prepare RNase I dilutions in nuclease-free water as follows:

  b. Add 10 µL of the appropriate **RNase I dilution** to each corresponding lysate (H, M, or L).
  c. Gently invert the tubes several times to mix, then briefly spin down.
  d. Immediately incubate the samples in a thermomixer at 37ºC for exactly 3 min with shaking at 1,100 rpm.
  e. Immediately place the samples on ice after incubation and keep it on ice for ≥ 3 min to stop RNA digestion.
  f. **Optional:** Load the samples onto Proteus mini clarification spin columns, centrifuge at 16,000 x *g* for 1 min at 4ºC and transfer the flow-throughs to new 1.5 mL tubes.

| RNase I | RNase I stock (μL) | Nuclease-free water (μL) |
| --- | --- | --- |
| High (H) | 6 | 24 |
| Medium (M) | 1 | 99 |
| Low (L) | 1 | 499 |

**Note:** For lysates from human HeLa and mouse P19 cells, a 1:100 – 1:150 dilution of RNase I typically yields RNA fragments in the desired size range. Optimal RNase I dilution must be empirically determined for other cell types or for lysates with different protein amounts and concentrations. Always perform RNase I optimization using lysates containing the same protein amount and concentration intended for the iCLIP3 library preparation experiment.

#### Immunoprecipitation

**Day 1 Timing: 2.5 h**

11. Cleanup of antibody-bead complexes.
  a. Briefly spin down the antibody–bead coupling mixture.
  b. Place the tube on a magnetic rack to collect the beads and discard the supernatant.
  c. Wash the beads twice with 750 µL Lysis buffer.
  d. Resuspend the beads for each sample in 100 µL Lysis buffer supplemented with 1X PIs.
  **Note**: For six samples, resuspend the beads in 630 µL Lysis buffer supplemented with 1X PIs.
12. Immunoprecipitation.
  a. Add 100 µL antibody-coupled beads from Step 11d to each RNase I-treated lysate from Step 10e (or Step 10f, if performed).
  b. Incubate the lysates with the beads on a rotating wheel for 2 h at 4ºC.
  **Note**: Adjust the incubation time according to the optimized immunoprecipitation conditions for the RBP of interest.
13. Immunoprecipitation cleanup.
  a. Briefly spin down the samples and place them on a magnetic rack to collect the beads.
  b. Discard the supernatant.
  c. Wash the beads twice with 800 µL High Salt buffer. Incubate the second wash for 5 min on a rotating wheel at 4ºC. After incubation, briefly spin down the tubes to collect beads from the tube lids before proceeding.
  d. Wash the beads twice with 800 µL Lysis buffer. During the second Lysis buffer wash, transfer the beads to new 1.5 mL tubes.
  e. Wash beads once with 800 µL PNK Wash buffer.
  f. Perform a final wash with 300 µL PNK Wash buffer.
  g. Keep the beads on ice before proceeding to the next step.

**Pause point**: The beads can be kept in the PNK Wash buffer on ice for 1–2 h.

#### 3′ RNA dephosphorylation

**Day 1 Timing: 45 min**

RNase I generates RNA fragments with 2′,3′-cyclic phosphates and 3′-phosphates, which must be converted to 3′-hydroxyl groups to enable efficient ligation of pCp-IR750 and DNA adapters. To achieve this, crosslinked RNA fragments are dephosphorylated on beads using T4 polynucleotide kinase (PNK). T4 PNK has 3′ phosphatase activity in the absence of ATP and in buffers at pH 6.0-6.5.

14. 3′ RNA dephosphorylation.
  a. Prepare the RNA Dephosphorylation buffer, mix thoroughly by pipetting and keep on ice.

  b. Place the beads from Step 13g on a magnetic rack and remove the PNK Wash buffer.
  c. Briefly spin down the tubes, return them to the magnetic rack and remove any residual buffer using a P20 pipette. **Note**: Process one sample at a time to avoid drying the beads.
  d. Resuspend the beads in 19 µL RNA Dephosphorylation buffer.
  e. Add 1 µL **T4 PNK** and mix thoroughly by pipetting.
  f. Incubate the samples for 20 min at 37ºC in a thermomixer with shaking at 1,200 rpm.
  **Note**: Place the sample in the thermomixer immediately after assembling the reaction to prevent bead sedimentation.
15. Cleanup of 3′ RNA dephosphorylation.
  a. Add 100 µL PNK Wash buffer without resuspending the beads.
  b. Place the tubes on a magnetic rack to collect the beads and remove the PNK Wash buffer.
  c. Wash the beads twice with 800 µL High Salt buffer. Incubate the second wash for 2 min on a rotating wheel at 4ºC. After incubation, briefly spin down the tubes to collect beads from the tube lids before proceeding.
  d. Wash the beads with 800 µL Lysis buffer. Transfer the beads to new 1.5 mL tubes during this wash.
  e. Wash the beads once with 800 µL PNK Wash buffer.
  f. Perform a final wash with 300 µL PNK Wash buffer.
16. Thoroughly resuspend the beads by pipetting and divide them into two tubes as follows:
  - Tube 1: 270 µL (90% beads)
  - Tube 2: 30 µL (10% beads)

| Reagent | Volume per reaction (μL) |
| --- | --- |
| Nuclease-free water | 14.5 |
| 5X T4 PNK buffer (pH 6.5) | 4 |
| RNasin | 0.5 |
| **Total** | **19** |

**Pause point**: Store the samples at 4ºC overnight.

#### 3’ RNA labeling with pCp-IR750

**Day 2 Timing: 1.5 h**

In this step, pCp-IR750 is ligated to 10% of the immunopurified RNA fragments that have undergone T4 PNK-mediated 3′-end repair (**Fig. 3A, B**). Labeling this subset of RNA fragments is sufficient to enable rapid and safe visualization of RBP–RNA complexes on a nitrocellulose membrane.

**Figure 3.**
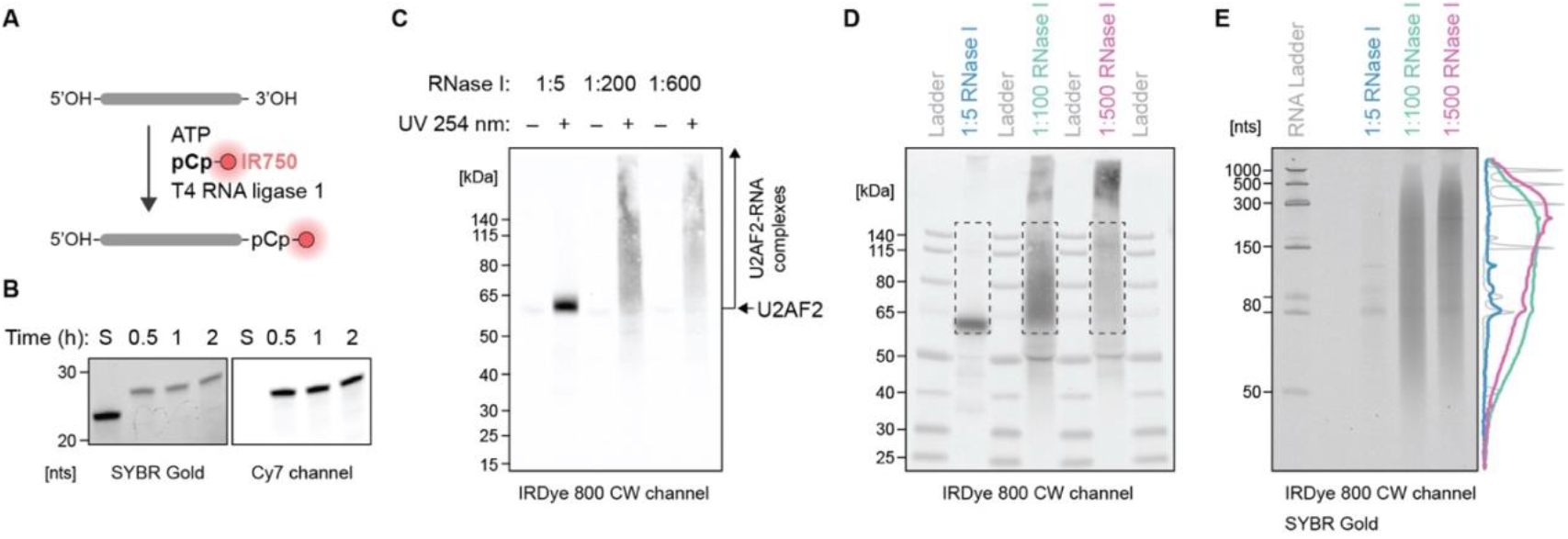
pCp-IR750 RNA labeling enables rapid and safe visualization of RNA–protein complexes. **(A)** Infrared 3′-end RNA labeling requires pCp-IR750, ATP, and T4 RNA ligase 1. **(B)** Efficiency of infrared 3′-end RNA labeling with pCp-IR750. A 15% TBE–urea gel shows attachment of pCp-IR750 to a 24-nucleotide (nt) RNA oligonucleotide. RNA was labeled with pCp-IR750 for 0.5, 1, and 2 h at 27ºC. Lane S contains unlabeled (input) RNA. The gel was stained with SYBR Gold and imaged in both SYBR Gold and Cy7 channels. **(C)** iCLIP3-based visualization of immunopurified protein–RNA complexes. A nitrocellulose membrane shows pCp-IR750-labeled RNA associated with immunopurified U2AF2–RNA complexes from UV-irradiated and non-irradiated samples treated with different RNase I dilutions (high, 1:5; medium, 1:200; low, 1:600). Only 10% of the immunoprecipitation (IP) material was subjected to pCp-IR750 RNA labeling. **(D)** Nitrocellulose membrane showing pCp-IR750-labeled RNA on 10% of the immunopurified U2AF2–RNA complexes from UV-irradiated samples treated with three RNase I dilutions (high, 1:5; medium, 1:100; low, 1:500). The remaining 90% of the IP material was unlabeled but run on the same gel. Dashed boxes indicate the membrane regions from which RNA was isolated. **(E)** RNA isolated from the membrane regions shown in (D) was labeled with pCp-IR750 and resolved on a 10% TBE–urea gel alongside a low-range single-stranded RNA ladder. The gel was imaged in SYBR Gold and IRDye 800CW channels.

17. 3′ RNA labeling with pCp-IR750.
  a. Prepare the RNA Labeling buffer, mix thoroughly by pipetting and keep at room temperature.

  b. Place Tube 2 (10% beads) on a magnetic rack to collect the beads and remove the PNK Wash buffer.
  c. Resuspend the beads in 19 µL RNA Labeling buffer.
  d. Add 1 µL **T4 RNA Ligase 1** (high concentration).
  e. Mix the reaction thoroughly by pipetting until the beads are evenly distributed. Pipette slowly to avoid bead retention in the pipette tip.
  f. Incubate the samples at 27ºC for 1 h in a thermomixer with shaking at 1,200 rpm.
  **Note**: Do not incubate the labeling reaction longer than 1 h to avoid protein labeling. Protect the samples from light using aluminum foil.
18. Cleanup of 3′ RNA labeling reaction.
  a. Add 100 µL PNK Wash buffer without resuspending the beads.
  b. Place the samples on a magnetic rack to collect the beads and remove the PNK Wash buffer.
  c. Wash the beads twice with 400 µL High Salt buffer.
  d. Wash the beads with 400 µL Lysis buffer. Transfer the beads to new 1.5 mL tubes during this wash.
  e. Wash the beads once with 400 µL PNK Wash buffer.
  f. Perform a final wash with 100 µL PNK Wash buffer.

| Reagent | Volume per reaction (μL) |
| --- | --- |
| Nuclease-free water | 7.75 |
| 10X RNA Ligase Reaction buffer | 2 |
| 10 mM ATP | 2 |
| 200 μM pCp-IR750 | 1 |
| RNasin | 0.25 |
| 50% PEG8000 | 6 |
| **Total** | **19** |

**Note:** Pipette the assembled labeling buffer slowly and thoroughly to ensure homogeneous distribution of all components. PEG8000 is highly viscous; avoid liquid retention in the pipette tip. Protect the buffer from light.

#### Elution of RBP–RNA complexes

**Day 2 Timing: 20 min**

Non-labeled (90%) and pCp-IR750-labeled (10%) RBP–RNA complexes are combined, washed, and eluted from the beads. Labeling a subset of RNA fragments with pCp-IR750 allows direct visualization of immunopurified RBP–RNA complexes on a nitrocellulose membrane. The remaining, non-labeled RNA (90%) is subsequently labeled with pCp-IR750 after RNA isolation from the nitrocellulose membrane.

19. Combine beads from Tube 1 and Tube 2.
  a. Transfer the beads from Tube 2 (100 µL) into Tube1 (270 µL).
  b. Wash the combined beads with 300 µL Last Wash buffer.
  c. Discard the supernatant.
20. Elution of RBP–RNA complexes.
  a. Collect the beads on a magnetic rack and discard the Last Wash buffer.
  b. Resuspend the beads for each sample in 20 µL 1X LDS NuPAGE Loading buffer supplemented with 50 mM DTT.
  c. Incubate the beads in a thermomixer at 70ºC for 5 min with shaking at 1,100 rpm.
  d. Briefly spin down the samples, collect the beads on a magnetic rack and transfer the eluates to new 1.5 mL tubes.

#### SDS-PAGE and nitrocellulose transfer of the RBP–RNA complexes

**Day 2 Timing: 3 h**

The eluted RBP–RNA complexes are resolved by denaturing SDS-PAGE and transferred onto a nitrocellulose membrane. Proteins efficiently bind nitrocellulose; therefore, RNA that is directly crosslinked to proteins will remain attached to the membrane via the RBP. In contrast, background or non-crosslinked RNA will not stably bind to the membrane.

21. SDS-PAGE of RBP–RNA complexes.
  a. Prepare 0.5 L 1X NuPAGE MOPS SDS Running buffer using nuclease-free water.
  b. Prepare the Prestained Protein Ruler (protein ladder) dilution for a single gel lane by mixing 2.5 µL protein ladder stock solution with 17.5 µL 1X LDS NuPAGE Loading buffer. **Note**: For this experiment, prepare PreStained Protein Ruler dilution for 6 gel lanes by mixing 15 µL protein ladder stock solution and 105 µL 1X LDS NuPAGE Loading buffer.
  c. Assemble a 12-well 4-12% Bis-Tris SDS gel in the XCell SureLock module according to the manufacturer’s instructions.
  d. Load 20 µL of each sample in the following order:

  e. Run the gel at 180 V for 50-60 min.
22. Transfer of RBP–RNA complexes to nitrocellulose membrane.
  a. Prepare 1X NuPAGE Transfer buffer from 20X stock using nuclease-free water and 20% (vol/vol) methanol or ethanol.
  b. Cut the nitrocellulose membrane and four pieces of Whatman paper.
  c. Carefully open the gel cassette.
  d. Assemble the transfer sandwich using the XCell II module according to the manufacturer’s instructions. **Note**: Pre-wet the sponges and remove any air bubbles using a roller.
  e. Perform the transfer for 1.5 h at 30 V.
  **Note**: For proteins smaller than 50 kDa, 1 h at 30 V is usually sufficient. For large proteins (>180 kDa), longer transfer times may be required.

| PL | UV-<br>H | UV-<br>M | UV-<br>L | PL | UV+<br>H | PL | UV+<br>M | PL | UV+<br>L | PL |  |
| --- | --- | --- | --- | --- | --- | --- | --- | --- | --- | --- | --- |
| 1 | 2 | 3 | 4 | 5 | 6 | 7 | 8 | 9 | 10 | 11 | 12 |

where PL: Protein Ladder, UV-: non-irradiated, UV+: UV-irradiated, H: high RNase I, M: medium RNase I, L: low RNase I.

#### Membrane imaging

**Day 2 Timing: 15 min**

During this step, immunopurified RBP–RNA complexes labeled with pCp-IR750 are visualized directly on the nitrocellulose membrane.

23. Visualization of immunopurified RBP–RNA complexes.
  a. Disassemble the transfer sandwich.
  b. Using forceps, place the membrane in a clean plastic tray containing 1X PBS.
  c. Place the membrane onto a thin, transparent plastic foil and insert it into the imager. **Note:** Do not place the membrane directly onto the transilluminator plate to avoid cross-contamination with nucleic acids.
  d. Image the membrane using the Cy7 or IRDye 800 CW channel to visualize the RBP– RNA complexes.
  e. Image the membrane in the colorimetric channel to visualize the protein ladder.
  f. Merge the two images into a single composite image and save.
  g. Place the membrane back into the plastic tray containing 1X PBS.
24. Inspect the image and identify the region of the membrane showing RNA signal that corresponds to the size range of the immunopurified RBP–RNA complexes (**Fig. 3C, D**).

#### Elution of RNA from the membrane with Proteinase K treatment

**Day 2 Timing: 1 h**

This step serves to determine the size range of the immunopurified RNA fragments. To release crosslinked RNA from the nitrocellulose membrane, the regions containing RBP– RNA complexes are excised from UV+ samples and subjected to digestion by Proteinase K, a non-specific protease. Both pCp-IR750-labeled (10%) and non-labeled (90%) RNA are released from the membrane during this step.

25. Membrane cutting.
  a. Place the membrane on a transparent, thick foil.
  b. Using sterile scalpels, cut the sample lane between two proteins ladders corresponding to the size of the immunopurified RBP–RNA complexes in UV+ samples (**Fig. 3D**).
  c. Transfer the excised membrane region to a 6 or 10-cm dish.
  d. Shred the membrane further into smaller pieces.
  e. Prepare 1.5 mL tubes with 150 µL Proteinase K buffer.
  f. Using needle tips, transfer the membrane pieces into the 1.5 mL tubes containing the Proteinase K buffer.
  **Note**: Ensure that all membrane pieces are fully submerged in the Proteinase K buffer.
26. Proteinase K treatment.
  a. Add 10 µL **Proteinase K** (20 mg/mL) to each sample.
  b. Incubate the samples in a thermomixer at 37ºC for 20 min with shaking at 1,000 rpm.
  c. Continue the incubation in a thermomixer at 50ºC for 20 min with shaking at 1,000 rpm.
  d. Briefly spin down the samples and transfer the supernatants (∼155 µL) into new 1.5 mL tubes.

#### RNA isolation

**Day 2 Timing: 15 min**

RNA released after Proteinase K treatment is purified using a silica column-based approach.

27. Proteinase K inactivation.
  a. Add 45 µL nuclease-free water to the sample from Step 26d.
  b. Add 1 µL 0.5 M phenylmethylsulfonylfluoride (PMSF) and briefly vortex.
  c. Briefly spin down the samples and incubate at room temperature for ≥ 3 min.
28. RNA isolation using RNA Clean and Concentrator-5 kit.
  a. Add 400 µL (2X volume) RNA Binding buffer and briefly vortex.
  b. Briefly spin the samples, add 700 µL 100% Isopropanol (3.5X starting volume) and briefly vortex.
  c. Incubate the samples for 15 min at room temperature on a rotating wheel.
  d. Briefly spin down the samples, transfer 650 µL of each sample into the Zymo-Spin IC column and centrifuge at 5,000 x g for 30 s at room temperature.
  e. Discard the flow-through from the collection tube and add the remaining sample to the same column.
  f. Centrifuge at 5,000 x g for 30 s at room temperature and discard the flow-through.
  g. Wash the columns with 400 µL RNA Prep buffer, centrifuge at 5,000 x g for 30 s at room temperature and discard the flow-through.
  h. Wash the columns with 500 µL RNA Wash buffer (with ethanol added), centrifuge at 5,000 x g for 30 s at room temperature and discard the flow-through.
  i. Wash the columns with 250 µL RNA Wash buffer (with ethanol added), centrifuge at 9,000 x g for 30 s at room temperature and discard the flow-through.
  j. Centrifuge at 9,000 x g for additional 2 min at room temperature and transfer the columns to clean 1.5 mL tubes, being careful to avoid contact between the wash buffer and the columns.
  k. Add 9.2 µL nuclease-free water to each column, incubate at 37ºC for 2 min and then centrifuge at 15,000 x g for 1 min at room temperature.
  l. Discard the column and freeze the eluted RNA at -80ºC.

**Pause point**: RNA can be stored at -80ºC until the next day.

#### RNA labeling

**Day 2 Timing: 2.5 h**

In this step, 90% of the previously unlabeled, immunopurified RNA is labeled with pCp-IR750. Labeling after RNA isolation preserves the IR750 signal and minimizes photobleaching.

29. 3′ RNA labeling with pCp-IR750.
  a. Set up the RNA labeling reaction as follows:

  b. Mix thoroughly by pipetting and incubate the ligation reaction in a thermomixer at 27ºC for 2 h with shaking at 1,000 rpm.
30. RNA isolation using RNA Clean and Concentrator-5 kit.
  a. Add 60 µL nuclease-free water to the ligation reaction.
  b. Add 160 µL RNA Binding buffer and briefly vortex.
  c. Briefly spin the samples, then add 280 µL isopropanol and briefly vortex.
  d. Incubate the samples on a rotating wheel at room temperature for 15 min, protected from light.
  e. Apply the samples to the Zymo-Spin IC columns, centrifuge at 5,000 x g for 30 s at room temperature and discard the flow-through.
  f. Wash the columns with 400 µL RNA Prep buffer, centrifuge at 5,000 x g for 30 s at room temperature and discard the flow-through.
  g. Wash the columns with 500 µL RNA Wash buffer (with ethanol added), centrifuge at 5,000 x g for 30 s at room temperature and discard the flow-through.
  h. Wash the columns with 250 µL RNA Wash buffer (with ethanol added), centrifuge at 9,000 x g for 30 s at room temperature and discard the flow-through.
  i. Centrifuge at 9,000 x g for additional 2 min at room temperature and transfer the columns to clean microcentrifuge tubes, being careful to avoid contact between the wash buffer and the columns.
  j. Add 7.2 µL nuclease-free water to each column, incubate at 37ºC for 2 min and then centrifuge at 15,000 x g for 1 min at room temperature.
  k. Discard the columns and keep the eluted RNA on ice until further use.

| Reagent | Volume per reaction (μL) |
| --- | --- |
| Isolated RNA | 8.8 |
| 10X RNA Ligase Reaction buffer | 2 |
| 20 mM ATP | 1 |
| 200 μM pCp-IR750 | 1 |
| RNasin | 0.2 |
| DMSO | 2 |
| 50% PEG8000 | 4 |
| T4 RNA ligase 1 (high concentration) | 1 |
| **Total** | **20** |

#### Denaturing RNA gel

**Day 3 Timing: 1.5 h**

In this step, pCp-IR750-labeled RNA is resolved on a denaturing 10% TBE-urea gel. This enables visualization of immunopurified RNA fragments and determination of their size range.

31. Separation of pCp-IR750 labeled RNA on a denaturing RNA gel.
  a. Prepare 7 µL Low Range Single-Stranded RNA (ssRNA) Ladder diluted 1:5 in nuclease-free water.
  b. Add 5 µL 2X TBE–Urea Sample buffer to each RNA sample from Step 30k and to the diluted ssRNA ladder.
  c. Heat the samples and the ssRNA ladder at 65ºC for 5 min prior to loading.
  d. Immediately place the samples and the ssRNA ladder on ice.
  e. Assemble a 10% TBE-urea gel in the XCell SureLock module and fill the chamber with 1X TBE Running buffer according to the manufacturer’s instructions.
  f. Use a P1000 pipette tip and flush any precipitated urea from the wells.
  g. Load 12 µL of each sample and the ssRNA ladder into the wells.
  h. Run the gel for 60 min at 200 V.
  i. Open the gel cassette and carefully remove the gel.
  j. Stain the gel for 5 min in 30 mL 1X TBE buffer containing 3 µL SYBR Gold.
  k. Visualize the pCp-IR750-labeled RNA using the Cy7 or IRDye 800 CW channel.
  l. Continue imaging the gel using the SYBR Gold channel to visualize the ssRNA ladder.
  m. Merge images from both channels to determine the size range of immunopurified RNA fragments based on the ssRNA ladder migration (**Fig. 3E**).

### EXPECTED OUTCOMES

Protocol 1 allows researchers to qualitatively assess whether the protein of interest directly binds RNA and to optimize RNase I-based RNA fragmentation. RNA labeling of the sample with high RNase I dilution should produce a strong pCp-IR750 signal in the UV+ sample, appearing as a single band corresponding to the molecular weight of the protein of interest, whereas the UV– sample should show minimal or ideally no infrared RNA signal. In UV+ samples treated with medium and low RNase I dilutions, the distinct RBP–RNA complex band gradually disperses into an upward smear, while the respective UV– samples remain devoid of infrared RNA signal (**Fig. 3C**). This shift in signal confirms that the visualized complexes contain RNA. For HeLa and mouse P19 cell lines, the optimal RNase I dilution lies within the 1:100–1:150 range in lysates containing 250 µg total protein at 0.5 µg/µL protein concentration, resulting in the majority of immunopurified RNA fragments in 50– 300 nt size range (**Fig. 3D, E**). The same RNase I treatment should not be applied to lysates with different protein or RNA amounts or concentrations. During RNase I optimization, all samples should be adjusted to match the sample with the lowest protein amount and concentration.

### TROUBLESHOOTING

#### Problem 1

No IP-grade antibody is available for the RBP of interest.

#### Potential solution

Use CRISPR-based technologies to insert a protein tag (e.g., GFP, HA) at the N- or C-terminus of the RBP. Then, perform IPs using commercially available antibodies or nanobodies specific to the inserted tag. Heterozygous tagging might be sufficient for an iCLIP3 experiment. Tagged proteins can also be transiently expressed from a transfected plasmid. However, transient expression should be carefully titrated as protein overexpression can lead to unspecific RNA binding of the protein and should hence be avoided.

#### Problem 2

Infrared RNA signal appears weak on the nitrocellulose membrane, even after long exposure (≥ 3 min) in UV+ samples, while UV– samples show no visible RNA signal.

#### Potential solution

The protein of interest may still bind RNA specifically, but the starting material could be limiting due to low protein expression or low RNA binding capacity. Increase the amount of starting material, such as the number of cells. Importantly, optimize RNase I treatment on the resulting lysates with higher protein amount and concentration or use multiple 250 µg lysate tubes at 0.5 mg/mL for RNase treatment and combine them prior to immunoprecipitation. In some cases, increasing the amount of antibody or using an alternative antibody can help.

#### Problem 3

Infrared RNA signal appears after longer exposure and looks the same in UV– and UV+ samples at high RNase I dilution. A distinct band does not disperse into an upward smear at medium and low RNase I dilution.

#### Potential solution

The protein of interest may interact weakly, transiently or not at all with RNA nucleobases. Consider increasing the UV 254 nm energy dose.

#### Problem 4

Multiple infrared bands are observed on the nitrocellulose membrane in the UV+ sample with high RNase I dilution.

#### Potential solution

Increase the stringency of the washing steps following the IP. Refer to the following publication for the recommended buffer recipes (Huppertz et al., 2014).

#### Problem 5

RNA signal is observed in the control samples where the specific antibody was omitted during the immunoprecipitation.

#### Potential solution

As for Problem 4, increase the stringency of the washing steps following the IP. Refer to the following publication for the recommended buffer recipes (Huppertz et al., 2014).

## PROTOCOL 2 iCLIP3 LIBRARY PREPARATION PROTOCOL

The following protocol can be used to prepare iCLIP3 sequencing libraries from UV+ samples using an antibody for the RBP of interest. We recommend including two control samples: A UV– sample using the antibody for the RBP of interest and a UV+ sample using non-immune IgG and/or omitting the antibody (bead-only).

### STEP-BY-STEP METHOD DETAILS

#### Cell culture and UV irradiation

##### Timing: variable, 2 days for the sample preparation

1. Cell seeding for the experiment.
  a. Grow HeLa cells in standard 1X DMEM medium supplemented with 10% FBS and 1X Penicillin-Streptomycin (Pen-Strep).
  b. One day before the experiment, seed HeLa cells in a 100 mm dish to achieve ∼85% confluency the following day.
2. PBS cell wash.
  a. Aspirate the culture medium from the dishes.
  b. Gently wash the cells with 6 mL ice-cold PBS.
  c. Remove the PBS and add 6 mL fresh ice-cold PBS to the dishes.
3. UV irradiation and cell harvesting.
  a. Place the cell dishes containing PBS on an ice tray covered with a thin layer of water.
  b. Remove the lids and irradiate the cells once with 254 nm UV light at 150 mJ/cm^2^.
  c. Gently scrape the cells using a cell lifter.
  d. Transfer the cell suspensions to 15 mL Falcon tubes.
  e. Pellet the cells by centrifugation at 300 x *g* for 5 min at 4ºC. **Note**: If available, use a swing-bucket rotor to ensure the cell pellets form at the bottom of the Falcon tubes.
  f. Carefully aspirate the PBS without disturbing the cell pellets.
  g. Snap-freeze the cell pellets using dry ice or liquid nitrogen and store at -80ºC.

**Pause point**: Frozen cell pellets can be stored at -80ºC long term until further use.

#### Antibody-bead coupling

**Day 1 Timing: 1-2 h**

4. Couple the antibody to magnetic Protein A or Protein G Dynabeads. **Note**: Verify the antibody’s binding preference for Protein A or Protein G Dynabeads according to the manufacturer’s recommendations.
  a. Aliquot 30 µL Protein A or Protein G Dynabeads per sample into a 1.5 mL tube. **Note**: Adjust the Dynabeads aliquot volume according to the number of samples.
  b. Wash the beads twice with 750 µL Lysis buffer.
  c. Resuspend the beads in 300 µL Lysis buffer and add 5 µg antibody per sample for the RBP of interest. **Note:** Adjust the amount of antibody based on the number of samples. Use 500 µL Lysis buffer for ≥ 6 samples. All washes in the protocol are performed by pipetting unless otherwise stated.
  d. Incubate the beads on a rotating wheel at room temperature for ≥ 1 h (until the lysates are ready for immunoprecipitation).

#### Cell lysis and protein quantification

**Day 1 Timing: 1 h**

5. Cell lysis.
  a. Thaw the frozen cell pellets on ice.
  b. Resuspend the cell pellets in 750 µL Lysis buffer supplemented with 1X PIs.
  c. Lyse the cells on ice for 10 min.
  d. **Optional:** Sonicate the lysates on ice at 10% amplitude using 5 cycles of 5 s pulses with 10 s pauses between pulses. **Note**: Sonication is recommended for nuclear proteins. If sonication is omitted, extend the lysis time to 20 min on ice.
  e. Transfer the lysates to new 1.5 mL tubes.
  f. Clarify the lysates by centrifugation at 16,000 x *g* for 10 min at 4ºC.
  g. Transfer the supernatants to fresh 1.5 mL tubes.
6. Determine the protein concentration of the cleared lysates using a BCA kit according to the manufacturer’s instructions.
7. Preparation the lysates for RNA fragmentation.
  a. Dilute the lysates to a final protein concentration of 0.5 mg/mL using Lysis buffer supplemented with 1X PIs.
  b. Transfer 500 µL of each diluted lysate (corresponding to 250 µg of total protein) into a new 1.5 mL tube and keep the samples on ice.

#### RNA fragmentation

**Day 1 Timing: 20 min**

The iCLIP3 technology relies on short-read sequencing. In this step, RNA in the lysate is partially digested with an appropriate concentration of RNase I to generate RNA fragments in the 50–300 nt size range. For lysates from human HeLa and mouse P19 cells, a 1:100– 1:150 dilution of RNase I typically yields RNA fragments in the desired size range. Optimal RNase I dilution must be empirically determined for other cell types and for lysates with different protein amounts and concentrations (see Protocol 1).

8. DNase treatment.
  a. Add 2 µL **TURBO DNase** to the diluted lysates from Step 7b.
  b. Gently invert the tubes several times to mix.
  c. Briefly spin down the samples and place on ice.
9. RNA fragmentation with RNase I.
  a. Prepare a low **RNase I dilution** in nuclease-free water as follows:

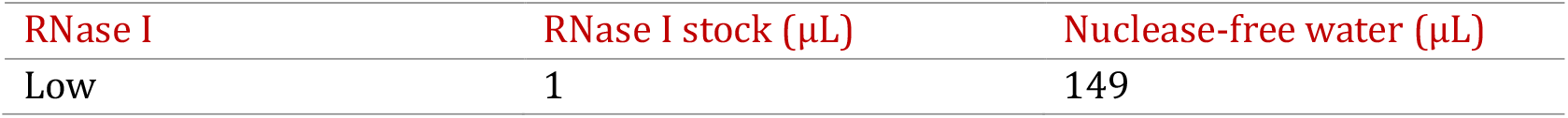 Add 10 µL low **RNase I dilution** to the lysate.
  b. Gently invert the tubes several times to mix, then briefly spin down.
  c. Immediately incubate the samples in a thermomixer at 37ºC for exactly 3 min with shaking at 1,100 rpm.
  d. Immediately place the samples on ice after incubation and keep them for ≥ 3 min to stop the RNA digestion. **Optional:** Load the samples onto Proteus mini clarification spin columns, centrifuge at 16,000 x *g* for 1 min at 4ºC and transfer the flow-throughs to new 1.5 mL tubes.

#### Immunoprecipitation

**Day 1 Timing: 2.5 h**

10. Cleanup of antibody-bead complexes.
  a. Briefly spin down the antibody-bead coupling mixture.
  b. Place the tubes on a magnetic rack to collect the beads and discard the supernatant.
  c. Wash the beads twice with 750 µL Lysis buffer.
  d. Resuspend the beads for each sample in 100 µL Lysis buffer supplemented with 1X PIs.
11. Immunoprecipitation.
  a. Add 100 µL antibody-coupled beads from Step 10d to the RNase I-treated lysates from Step 9e (or Step 9f, if performed).
  b. Incubate the lysates with the beads on a rotating wheel for 2 h at 4ºC. **Note**: Adjust the incubation time according to the optimized immunoprecipitation conditions for the RBP of interest.
12. Immunoprecipitation cleanup.
  a. Briefly spin down the samples and place them on a magnetic rack to collect the beads.
  b. Wash the beads twice with 800 µL High Salt buffer. Incubate the second wash on a rotating wheel for 5 min at 4ºC. After incubation, briefly spin down the samples to collect beads from the tube lid before proceeding.
  c. Wash the beads twice with 800 µL Lysis buffer. During the second Lysis buffer wash, transfer the beads to a new 1.5 mL tube.
  d. Wash the beads once with 800 µL PNK Wash buffer.
  e. Perform a final wash with 300 µL PNK Wash buffer.
  f. Keep the beads on ice until proceeding to the next step.

**Pause point**: The beads can be kept in the PNK Wash buffer on ice for 1–2 h.

#### 3′ RNA dephosphorylation

**Day 1 Timing: 45 min**

RNase I generates RNA fragments with 2′,3′-cyclic phosphates and 3′-phosphates, which must be converted to 3′-hydroxyl groups to enable efficient ligation of pCp-IR750 and DNA adapters. To achieve this, crosslinked RNA fragments are dephosphorylated on beads using T4 polynucleotide kinase (PNK).

13. 3′ RNA dephosphorylation.
  a. Prepare the RNA Dephosphorylation buffer, mix thoroughly by pipetting and keep on ice.

  b. Place the samples from Step 12f on a magnetic rack to collect the beads and remove the PNK Wash buffer.
  c. Briefly spin down the tubes, return it quickly to the magnetic rack, and remove any residual buffer using a P20 pipette. **Note**: Process one sample at a time to avoid drying the beads.
  d. Resuspend the beads in 19 µL RNA Dephosphorylation buffer.
  e. Add 1 µL **T4 PNK** and mix thoroughly by pipetting.
  f. Incubate the samples for 20 min at 37ºC in a thermomixer shaking at 1,200 rpm. **Note**: Place the samples in the thermomixer immediately after assembling the reaction to prevent bead sedimentation.
14. Cleanup of 3′ RNA dephosphorylation.
  a. Add 100 µL PNK Wash buffer without resuspending the beads.
  b. Place the samples on a magnetic rack to collect the beads and remove the PNK Wash buffer.
  c. Wash the beads twice with 800 µL High Salt buffer. Incubate the second wash for 2 min on a rotating wheel at 4ºC. After incubation, briefly spin down the samples to collect beads from the tube lid before proceeding.
  d. Wash the beads with 800 µL Lysis buffer. Transfer the beads to a new 1.5 mL tube during this wash.
  e. Wash the beads once with 800 µL PNK Wash buffer.
  f. Perform a final wash with 300 µL PNK Wash buffer.
  g. Thoroughly resuspend the beads by pipetting and divide each sample into two 1.5 mL tubes as follows:
    - Tube 1: 270 µL (90% beads)
    - Tube 2: 30 µL (10% beads)
  h. Store Tube 2 (10% beads) at 4ºC overnight and keep Tube 1 (90% beads) on ice.

| Reagent | Volume per reaction (μL) |
| --- | --- |
| Nuclease-free water | 14.5 |
| 5X T4 PNK buffer (pH 6.5) | 4 |
| RNasin | 0.5 |
| Total | 19 |

#### 3′ L7 linker ligation

**Day 1–2 Timing: 1 h**

The 5′-preadenylated L7 linker, a DNA adapter, is ligated on beads to 90% of the immunopurified RNA fragments with T4 PNK-repaired ends. Ligation of the L7 linker is essential for converting RBP-bound RNA into iCLIP3 sequencing libraries.

15. 3′ L7 linker ligation.
  a. Prepare the RNA Ligation buffer, mix well by pipetting and keep it at room temperature:

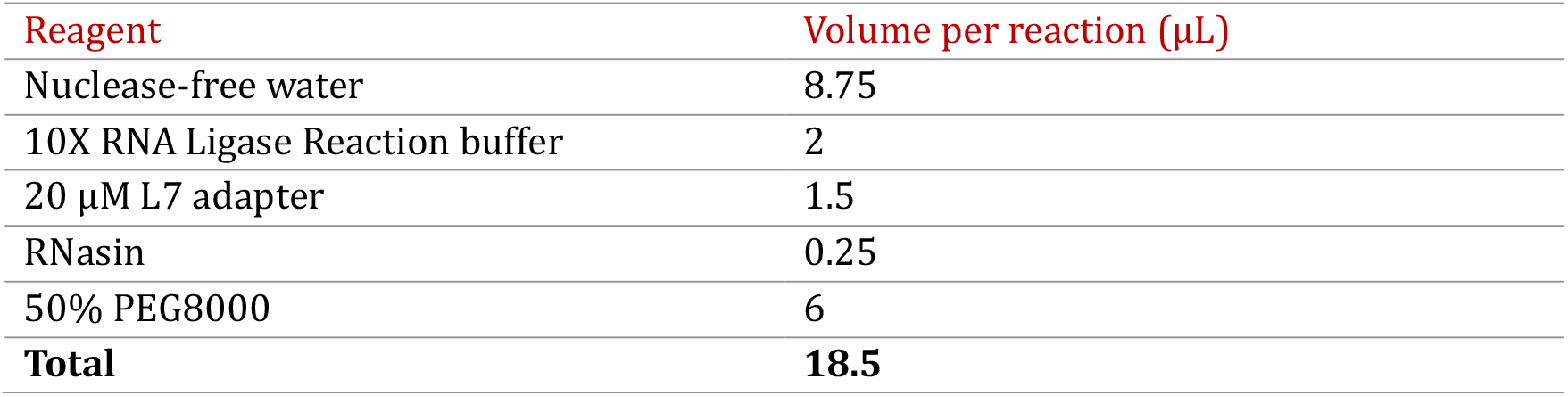 **Note:** Pipette the assembled RNA Ligation buffer slowly and thoroughly to ensure homogeneous distribution of all components. PEG8000 is highly viscous; avoid liquid retention in the pipette tip.
  b. Place each Tube 1 from Step 14e on a magnetic rack to collect the beads and remove the PNK Wash buffer.
  c. Briefly spin down the tubes, return them quickly to the magnetic rack, and remove any residual buffer using a P20 pipette. **Note**: Process one sample at a time to avoid drying the beads.
  d. Resuspend the beads in 18.5 µL RNA Ligation buffer.
  e. Add 1.5 µL **T4 RNA Ligase 1** (high concentration).
  f. Mix the reactions thoroughly by pipetting until the beads are evenly distributed. Pipette slowly to avoid bead retention in the pipette tip.
  g. Incubate the ligation reactions at 16ºC overnight in a thermomixer with shaking at 1,200 rpm.
16. Cleanup of 3′ L7 linker ligation (**Day 2**).
  a. Add 100 µL PNK Wash buffer without resuspending the beads.
  b. Place the samples on a magnetic rack to collect the beads and remove the PNK Wash buffer.
  c. Wash the beads twice with 800 µL High Salt buffer. Incubate the second wash for 2 min on a rotating wheel at 4ºC. After incubation, briefly spin down the samples to collect beads from the tube lid before proceeding.
  d. Wash the beads with 800 µL Lysis buffer. Transfer the beads to a new 1.5 mL tube during this wash.
  e. Wash the beads twice with 800 µL PNK Wash buffer.
  f. Leave the beads on ice in the PNK Wash buffer.

#### 3′ RNA labeling with pCp-IR750

**Day 2 Timing: 1.5 h**

pCp-IR750 is ligated on beads to 10% of the immunopurified RNA fragments with T4 PNK-repaired 3′ ends. pCp-IR750 RNA labeling enables rapid and safe visualization of RBP-RNA complexes on a nitrocellulose membrane.

17. 3′ RNA labeling with pCp-IR750.
  b. Prepare the RNA Labeling buffer, mix thoroughly by pipetting and keep at room temperature.

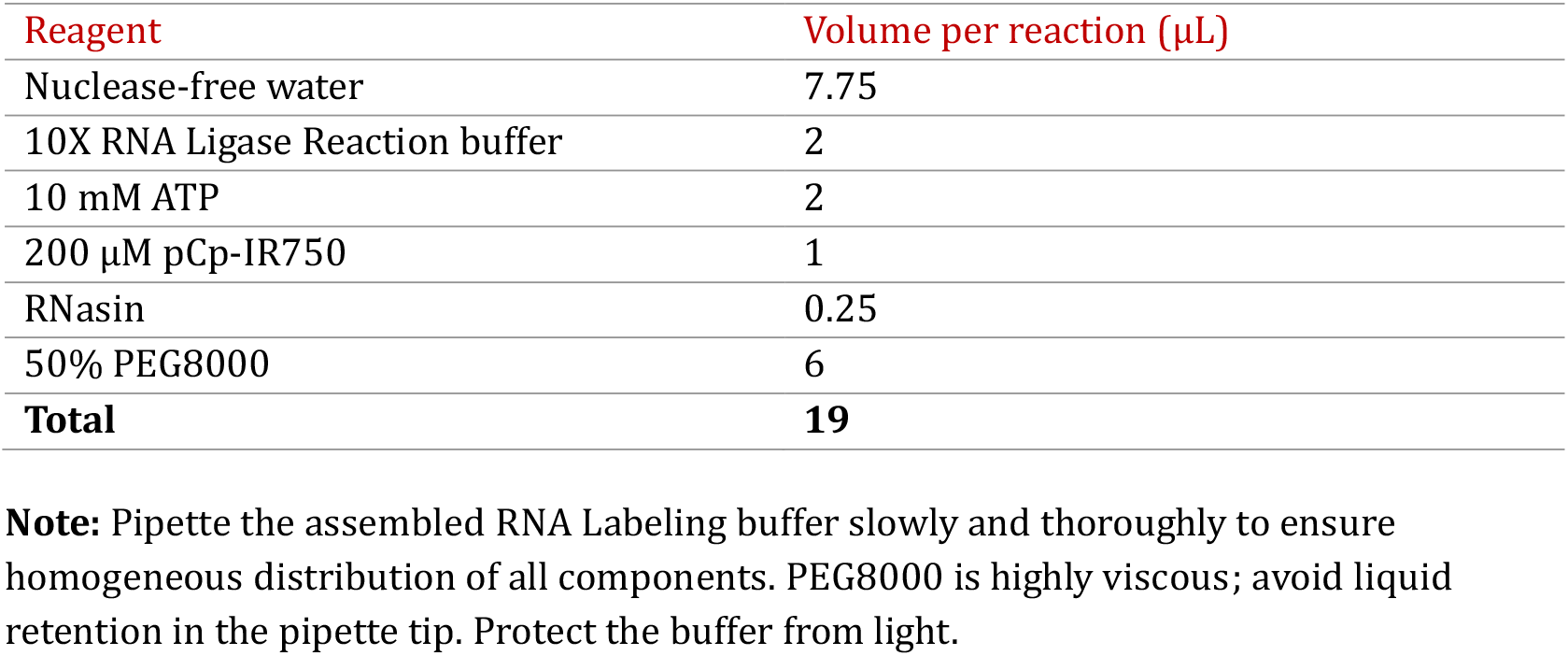
  c. Place each Tube 2 (10% beads) on a magnetic rack to collect the beads and remove the PNK Wash buffer.
  d. Resuspend the beads in 19 µL RNA Labeling buffer.
  e. Add 1 µL **T4 RNA Ligase 1** (high concentration).
  f. Mix the reactions thoroughly by pipetting until the beads are evenly distributed. Pipette slowly to avoid bead retention in the pipette tip.
  g. Incubate the reactions at 27ºC for 1 h in a thermomixer with shaking at 1,200 rpm. **Note**: Protect the samples from light using aluminum foil.
18. Cleanup of 3′ RNA labeling reaction.
  a. Add 100 µL PNK Wash buffer without resuspending the beads.
  b. Place the samples on a magnetic rack to collect the beads and remove the PNK Wash buffer.
  c. Wash the beads twice with 400 µL High Salt buffer.
  d. Wash the beads with 400 µL Lysis buffer. Transfer the beads to a new 1.5 mL tube during this wash.
  e. Wash the beads once with 400 µL PNK Wash buffer.
  f. Perform a final wash with 100 µL PNK Wash buffer.

#### Elution of RBP–RNA complexes

**Day 2 Timing: 20 min**

L7 linker-ligated and pCp-IR750-labeled RBP-RNA complexes are combined, washed and eluted from the beads. While pCp-IR750 RNA labeling allows visualization of the immunopurified RBP–RNA complexes, L7 linker ligation enables conversion of the RNA into iCLIP3 sequencing libraries.

19. For each sample, combine beads from Tube 1 and Tube 2.
  a. Transfer the beads from Tube 2 (100 µL) into Tube1 (800 µL).
  b. Wash the combined beads with 300 µL Last Wash buffer.
20. Elution of RBP-RNA complexes.
  a. Collect the beads on a magnetic rack and discard the Last Wash buffer.
  b. Resuspend the beads in 20 µL 1X LDS NuPAGE Loading buffer supplemented with 50 mM DTT.
  c. Incubate the beads in a thermomixer at 70ºC for 5 min with shaking at 1,100 rpm.
  d. Briefly spin down the samples, collect the beads on a magnetic rack, and transfer the eluates to new 1.5 mL tubes.

#### SDS-PAGE and nitrocellulose transfer of RBP–RNA complexes

**Day 2 Timing: 3 h**

The eluted RBP-RNA complexes are resolved by denaturing SDS-PAGE and transferred onto a nitrocellulose membrane. Proteins efficiently bind nitrocellulose; therefore, RNA that is directly crosslinked to proteins will remain attached to the membrane via the RBP. In contrast, background or non-crosslinked RNA will not stably bind to the membrane.

21. SDS-PAGE of RBP-RNA complexes.
  a. Prepare 0.5 L 1X NuPAGE MOPS SDS Running buffer using nuclease-free water.
  b. Prepare the Prestained Protein Ruler (protein ladder) dilution for two gel lanes per sample by mixing 6.25 µL protein ladder stock solution with 43.75 µL 1X LDS NuPAGE Loading buffer (includes 2.5x surplus).
  c. Assemble a 12-well 4-12% Bis-Tris SDS gel in the XCell SureLock module according to the manufacturer’s instructions.
  d. Load 20 µL protein ladder dilution in two gel lanes, leaving an empty gel lane between them.
  e. Load 20 µL sample from Step 20d in the empty lane between the protein ladder lanes.
  f. Run the gel at 180 V for 50-60 min.
22. Transfer of RBP–RNA complexes to a nitrocellulose membrane.
  a. Prepare 1X NuPAGE Transfer buffer with 20X buffer stock using nuclease-free water and 20% (vol/vol) methanol or ethanol.
  b. Cut the nitrocellulose membrane and four pieces of Whatman paper.
  c. Carefully open the gel cassette.
  d. Assemble the transfer sandwich using the XCell II module according to the manufacturer’s instructions. **Note**: Pre-wet the sponges and remove any air bubbles using a roller.
  e. Perform the transfer in 1X NuPAGE Transfer buffer for 1.5 h at 30 V. **Note**: For proteins smaller than 50 kDa, transfer for 1 h at 30 V. For large proteins (> 180 kDa), longer transfer times may be required.

#### Membrane imaging

**Day 2 Timing: 15 min**

In this step, immunopurified RBP-RNA complexes labeled with pCp-IR750 are visualized directly on the nitrocellulose membrane.

23. Visualization of immunopurified RBP–RNA complexes.
  a. Disassemble the transfer sandwich.
  b. Using forceps, place the membrane in a clean plastic tray containing 1X PBS.
  c. Place the membrane onto a thin, transparent plastic foil and insert it into the imager. **Note:** Do not place the membrane directly onto the transilluminator plate to avoid contamination from nucleic acids previously imaged.
  d. Image the membrane using the Cy7 or IRDye 800 CW channel to visualize the RBP– RNA complexes.
  e. Image the membrane using the colorimetric channel to visualize the protein ladder.
  f. Merge the two images into a single composite image and save.
  g. Return the membrane to the plastic tray containing 1X PBS.
24. Inspect the image and identify the region of the membrane showing RNA signal that corresponds to the size range of immunopurified RBP–RNA complexes (**Fig. 3C, D, 5A**).

#### Elution of RNA from the membrane with Proteinase K treatment

**Day 2 Timing: 1 h**

Nitrocellulose regions containing RBP-RNA complexes are excised and subjected to Proteinase K treatment. Proteinase K is a non-specific protease that liberates RNA from the membrane by digesting the protein. Both pCp-IR750-labeled (10%) and L7 linker-ligated (90%) RNA are released from the membrane by Proteinase K.

25. Membrane cutting.
  a. Place the membrane on a transparent, thick foil.
  b. Using sterile scalpels, cut the sample lane between two proteins ladders corresponding to the size of the immunopurified RBP–RNA complexes (**Fig. 3D, 5A**).
  c. Transfer the excised membrane region to a clean 6-cm or 10-cm dish.
  d. Shred the membrane further into smaller pieces.
  e. Prepare a 1.5 mL tube containing 150 µL Proteinase K buffer.
  f. Using needle tips, transfer the membrane pieces into the 1.5 mL tube containing the Proteinase K buffer. **Note**: Ensure that all membrane pieces are fully submerged in Proteinase K buffer.
26. Proteinase K treatment.
  a. Add 10 µL **Proteinase K** (20 mg/mL) to each sample.
  b. Incubate the samples in a thermomixer at 37ºC for 20 min with shaking at 1,000 rpm.
  c. Continue the incubation in a thermomixer at 50ºC for 20 min with shaking at 1,000 rpm.
  d. Briefly spin down the samples and transfer the supernatants (∼155 µL) to new 1.5 mL tubes.

#### RNA isolation

**Day 2 Timing: 15 min**

RNA released after Proteinase K treatment is purified using a silica column-based approach.

27. Proteinase K inactivation.
  a. Add 45 µL nuclease-free water to the samples from Step 26d.
  b. Add 1 µL 0.5 M PMSF and briefly vortex.
  c. Briefly spin down the samples and incubate at room temperature for ≥ 3 min.
28. RNA isolation using RNA Clean and Concentrator-5 kit.
  a. Add 400 µL (2 volumes) RNA Binding buffer and briefly vortex.
  b. Briefly spin the samples, add 700 µL 100% isopropanol (3.5X starting volume) and briefly vortex.
  c. Incubate the samples for 15 min at room temperature on a rotating wheel.
  d. Briefly spin down the samples, transfer 650 µL into the Zymo-Spin IC column and centrifuge at 5,000 x g for 30 s at room temperature.
  e. Discard the flow-through from each collection tube and add the remaining sample to the same column.
  f. Centrifuge at 5,000 x g for 30 s at room temperature and discard the flow-throughs.
  g. Wash the columns with 400 µL RNA Prep buffer, centrifuge at 5,000 x g for 30 s at room temperature and discard the flow-throughs.
  h. Wash the columns with 500 µL RNA Wash buffer (with ethanol added), centrifuge at 5,000 x g for 30 s at room temperature and discard the flow-throughs.
  i. Wash the columns with 250 µL RNA Wash buffer (with ethanol added), centrifuge at 9,000 x g for 30 s at room temperature and discard the flow-throughs.
  j. Centrifuge at 9,000 x g for additional 2 min at room temperature and transfer the columns to clean 1.5 mL tubes, being careful to avoid contact between the wash buffer and the column.
  k. Add 10.3 µL nuclease-free water to the columns, incubate at 37ºC for 2 min and then centrifuge at 15,000 x g for 1 min at room temperature.
  l. Discard the columns and store the eluted RNA at -80ºC.

**Pause point**: RNA can be stored at -80ºC until the next day.

#### Reverse transcription

**Day 3 Timing: 1.5 h**

In this step, eluted RNA is converted into cDNA using reverse transcriptase. The RT primer anneals to the 3′ end of the L7 linker, and reverse transcriptase synthesizes a cDNA copy. Proteinase K leaves a small peptide at the crosslinked nucleotide, which causes premature termination of reverse transcription and generates truncated cDNA.

29. cDNA synthesis.
  a. Thaw the isolated RNA on ice and transfer 10 µL to a PCR tube.
  b. Prepare the following master mix:

  c. Add 2 µL master mix to 10 µL isolated RNA and mix well.
  d. Incubate the mixture in a thermocycler with the lid heated to the maximum temperature (usually 105ºC) using the following program:

| Reagent | Volume per reaction ($\mu\text{L}$) |
| --- | --- |
| dNTP mix (10 mM each) | 1 |
| iCLIP3 RT primer (2 $\mu\text{M}$) | 1 |
| **Total** | **2** |

  e. Prepare the following RT mix, pipette thoroughly and keep on ice:

| Steps | Temperature | Time | Cycle number |
| --- | --- | --- | --- |
| Denaturation | 70°C | 5 min | 1 |
| Hybridization | 25°C | ∞ | 1 |

  f. Add 8 µL pre-assembled RT mix to each sample recovered from the thermocycler and mix well by pipetting.
  g. Return the samples back to the thermocycler and run the following program with the lid heated to the maximum temperature (usually 105ºC):

e. Prepare the following RT mix, pipette thoroughly and keep on ice:
| Reagent | Volume per reaction (μL) |
| --- | --- |
| Nuclease-free water | 2 |
| 5X First Strand buffer | 4 |
| 0.1 M DTT | 1 |
| RNasin | 0.5 |
| SuperScript III | 0.5 |
| **Total** | **8** |

| Steps | Temperature | Time | Cycle number |
| --- | --- | --- | --- |
| Initial incubation | 25°C | 5 min | 1 |
| cDNA synthesis | 42°C | 20 min | 1 |
|  | 50°C | 40 min |  |
| Inactivation | 85°C | 5 min | 1 |
| Hold | 16°C | ∞ | 1 |

#### RNA degradation

**Day 3 Timing: 30 min**

In this step, the RNA is hydrolyzed under alkaline conditions and elevated temperature to remove it from the samples.

30. RNA hydrolysis.
  a. Add 1.65 µL 1 M NaOH to each RT reaction and mix well.
  b. Place the samples back to the thermocycler and run the following program with the lid heated to the maximum temperature (usually 105ºC):

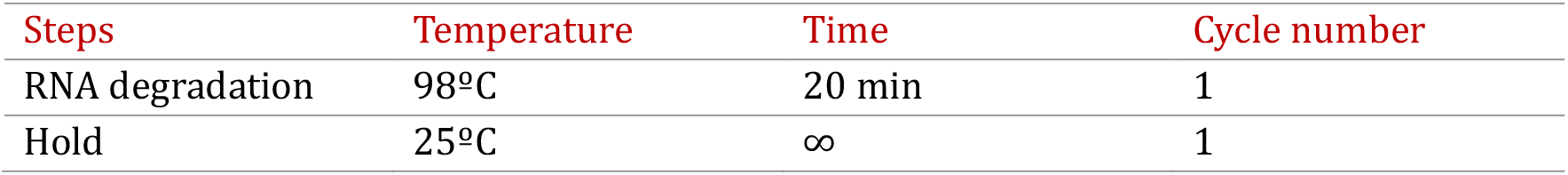
31. Neutralization.
  a. Add 20 µL 1 M HEPES to each sample recovered from the thermocycler and mix well.
  b. Transfer each sample to a non-stick 1.5 mL tube.

#### Isolation of truncated cDNA with MyOne Silane beads

**Day 3 Timing: 30 min**

In this step, the truncated cDNA is isolated using MyOne Silane beads.

32. Preparation of MyOne Silane beads for isolation of truncated cDNA.
  a. Aliquot 10 µL MyOne Silane beads per sample from the original stock into a 1.5 mL tube.
  b. Wash the beads once with 750 µL RLT buffer.
  c. Resuspend the beads in 125 µL RLT buffer per sample.
33. Isolation of truncated cDNA with MyOne Silane beads.
  a. Add 125 µL beads in RLT buffer to the samples from Step 31b and briefly mix by pipetting.
  b. Add 150 µL 100% ethanol to the samples and mix well by pipetting.
  c. Incubate the mixture at room temperature for 5 min.
  d. Pipette the mixture again and incubate at room temperature for additional 5 min.
  e. Place the samples on a magnetic rack to collect the beads and discard the supernatant.
  f. Wash the beads with 800 µL 80% ethanol by resuspending them.
  g. Transfer the beads in 80% ethanol to a new 1.5 mL non-stick tube.
  h. Place the samples on a magnetic rack to collect the beads and discard the supernatant.
  i. Wash the beads twice with 80% ethanol without resuspending.
  j. After the last wash, slowly lift the tube up on the magnetic rack to collect the beads at the bottom in a single spot.
  k. Discard the supernatant and air-dry the beads. **Note**: Do not over-dry the beads.
  l. Resuspend the beads in 5.3 µL nuclease-free water.

#### Second adapter ligation to 3′ ends of truncated cDNA

**Day 3 Timing: 20 min**

In this step, a 5′-phosphorylated DNA adapter containing unique molecular identifiers (5′UMIs) is ligated to the 3′ end of the cDNA at the truncation site.

34. Second adapter ligation.
  a. Prepare the following master mix:

  b. Add 2 µL master mix to the beads from Step 33l.
  c. Heat the samples at 75ºC for 2 min, then place on ice for 30 s and allow them to equilibrate to room temperature.
  d. Prepare the following additional master mix:

| Reagent | Volume per reaction (μL) |
| --- | --- |
| 100% DMSO | 1 |
| 20 μM iCLIP3 second adapter | 1 |
| **Total** | **2** |

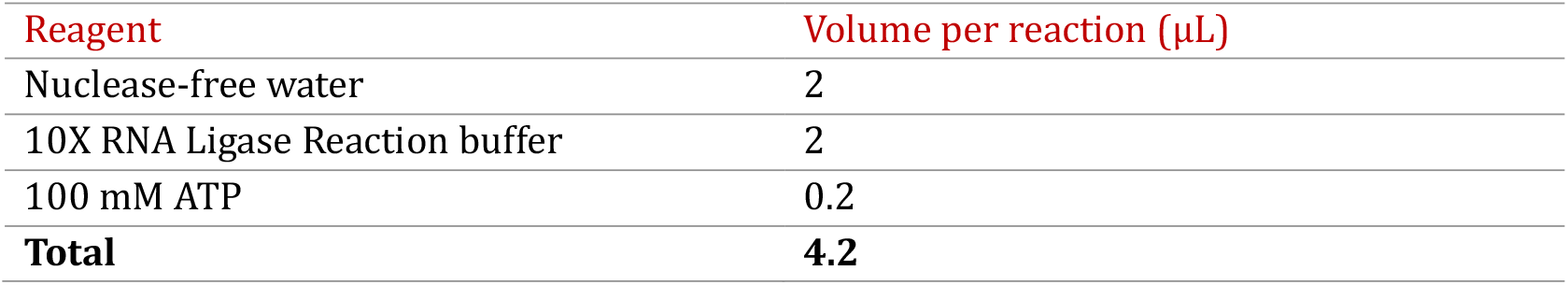
  e. Add 4.2 µL additional master mix to the beads from Step 34c.
  f. Add 8 µL 50% PEG8000 and 1.5 µL **T4 RNA Ligase 1** (high concentration).
  g. Mix the assembled ligation reactions thoroughly by pipetting. **Note:** Pipette the assembled ligation reactions slowly and thoroughly to ensure homogeneous distribution of all components. PEG8000 is highly viscous; avoid liquid and bead retention in the pipette tip.
  h. Incubate the ligation reactions in a thermomixer at 25ºC overnight with shaking at 1,200 rpm.

| Reagent | Volume per reaction (μL) |
| --- | --- |
| Nuclease-free water | 2 |
| 10X RNA Ligase Reaction buffer | 2 |
| 100 mM ATP | 0.2 |
| **Total** | **4.2** |

#### Isolation of the final cDNA with MyOne Silane beads

**Day 4 Timing: 30 min**

In this step, the final cDNA containing both adapters is recovered from the second adapter ligation reaction using MyOne Silane beads.

35. Preparation of MyOne Silane beads for isolation of final cDNA.
  a. Aliquot 5 µL MyOne Silane beads per sample from the original stock in a 1.5 mL tube.
  b. Wash the beads once with 750 µL RLT buffer.
  c. Resuspend the beads in 60 µL RLT buffer per sample.
36. Isolation of the final cDNA with MyOne Silane beads.
  a. Add 60 µL beads in RLT buffer to the ligation reaction from Step 34h and briefly mix by pipetting.
  b. Add 60 µL 100% ethanol to the samples and mix well by pipetting.
  c. Incubate the mixture at room temperature for 5 min.
  d. Pipette the mixture again and incubate at room temperature for additional 5 min.
  e. Place the samples on a magnetic rack to collect the beads and discard the supernatant.
  f. Wash the beads with 800 µL 80% ethanol by resuspending them.
  g. Transfer the beads in 80% ethanol to a new 1.5 mL non-stick tube.
  h. Place the samples on a magnetic rack to collect the beads and discard the supernatant.
  i. Wash the beads twice with 80% ethanol without resuspending.
  j. After the last wash, slowly lift the tube up on the magnetic rack to collect the beads at the bottom in a single spot.
  k. Discard the supernatants and airdry the beads. **Note**: Do not over-dry the beads.
  l. Resuspend the beads in 15 µL nuclease-free water and incubate at room temperature for 5 min.
  m. Place the samples on a magnetic rack and transfer each supernatant (14.7µL) to a new 1.5 mL tube.

**Pause point**: The isolated final cDNA can be stored at -20ºC long term.

#### Short-primer PCR (final cDNA pre-amplification)

**Day 4 Timing: 30 min**

In this step, the final cDNA is amplified with short primers using a low number of PCR cycles. The short-primer PCR generates a short iCLIP3 library, which is not yet ready for sequencing. The short iCLIP3 library can subsequently be size-selected to remove contaminating adapter dimers.

37. Short-primer PCR.
  a. Assemble the following PCR reaction:

  b. Run the PCR in a thermocycler using the following PCR program:

| Reagent | Volume per reaction (μL) |
| --- | --- |
| Final cDNA from Step 36m | 14.7 |
| Primer mix of P5s and P7s (50 μM each) | 0.3 |
| 2X Phusion HF PCR MasterMix | 15 |
| **Total** | **30** |

**Note:** It is essential to keep the PCR reaction volume exactly at 30 μL.

b. Run the PCR in a thermocycler using the following PCR program:
| Steps | Temperature | Time | Cycle number |
| --- | --- | --- | --- |
| Denaturation 1 | 98°C | 30 s | 1 |
| Denaturation 2 | 98°C | 10 s | 5-7 |
| Annealing | 65°C | 30 s |  |
| Extension | 72°C | 30 s |  |
| Final extension | 72°C | 3 min | 1 |
| Hold | 16°C | ∞ | 1 |

**Note:** For abundant RBPs with known RBDs, perform 5 PCR cycles. For unknown RBPs, perform 6 PCR cycles. For proteins that bind RNA weakly or transiently, 7 PCR cycles may be required.

#### Short iCLIP3 library size selection with ProNex beads

**Day 4 Timing: 30 min**

In this step, ProNex beads are used to size-select the short iCLIP3 library. This step removes adapter dimers (∼50 bp) and recovers the library ≥ 75 bp (insert size: ≥ 25 bp), with the most efficient recovery for library ≥ 100 bp (insert size: 50 bp). Ultra Low Range (ULR) DNA ladder is used to evaluate ProNex-based size selection.

38. Equilibrate ProNex beads and ProNex Wash buffer (with ethanol added) at room temperature for 30 min prior to use.
39. Short iCLIP3 library size selection.
  a. Set up the size selection ULR ladder control.

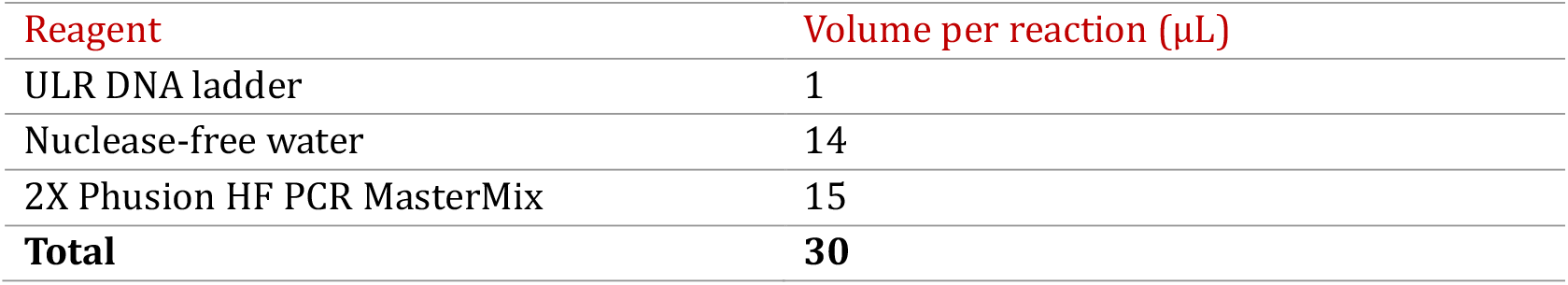
  b. Set up the reference ULR ladder and keep at 4ºC until further use.

| Reagent | Volume per reaction (μL) |
| --- | --- |
| ULR DNA ladder | 1 |
| Nuclease-free water | 14 |
| 2X Phusion HF PCR MasterMix | 15 |
| **Total** | **30** |

  c. Vortex the ProNex bead stock solution and mix thoroughly by pipetting.
  d. Add 2.95 volumes of ProNex beads (88.5 µL) to the PCR reaction and to the size selection ULR ladder control from Step39a.
  e. Mix well by pipetting up and down 8–10 times.
  f. Incubate the mixtures at room temperature for 10 min.
  g. Place the tubes on a magnetic rack to collect the beads.
  h. Discard the supernatants.
  i. Add 150 µL ProNex Wash buffer without disturbing the beads and incubate 15 s.
  j. Repeat steps h and i.
  k. Discard the supernatants and air-dry the beads until bead cracking becomes apparent. **Note**: Beads can be dried for 1–2 min at 37ºC. Do not over-dry the beads.
  l. Remove the tubes from the magnetic rack and resuspend the beads in 23 µL nuclease-free water.
  m. Incubate the samples for 5 min at room temperature.
  n. Place the tubes on a magnetic rack to collect the beads.
  o. Transfer the eluted short iCLIP library and size selected ULR ladder control to clean 1.5 mL tubes.

- b. Set up the reference ULR ladder and keep at 4°C until further use.
| Reagent | Volume per reaction (μL) |
| --- | --- |
| ULR DNA ladder | 1 |
| Nuclease-free water | 22 |
| **Total** | **23** |

**Pause point**: The eluted short iCLIP3 library and the size selected ULR ladder control can be stored at -20ºC long term.

#### Evaluation of ProNex size selection of short iCLIP3 library

**Day 4 Timing: 20 min**

This step evaluates the efficiency of ProNex size selection using the reference ULR ladder and size-selected ULR ladder control.

40. Evaluation of ProNex size selection using the TapeStation.
  a. Equilibrate TapeStation reagents at room temperature for 20–30 min prior to use.
  b. Load 2 µL reference ULR ladder from Step 39b and 2 µl size selected ULR ladder control from Step 39o onto the TapeStation using High Sensitivity D1000 ScreenTape.
  c. Assess the recovery of the 50 bp, 75 bp and 100 bp bands in the size selected ULR ladder control relative to the reference ULR ladder. **Note**: Successful size selection is indicated by significant removal of the 50-bp band, ∼50% recovery of the 75-bp band, and ∼100% recovery of the 100-bp band (**Fig. 4A**). Additionally, the ratio of the 75-bp to 50-bp band intensities in the size selected ULR ladder control should be ≥ 2.5.

**Figure 4.**
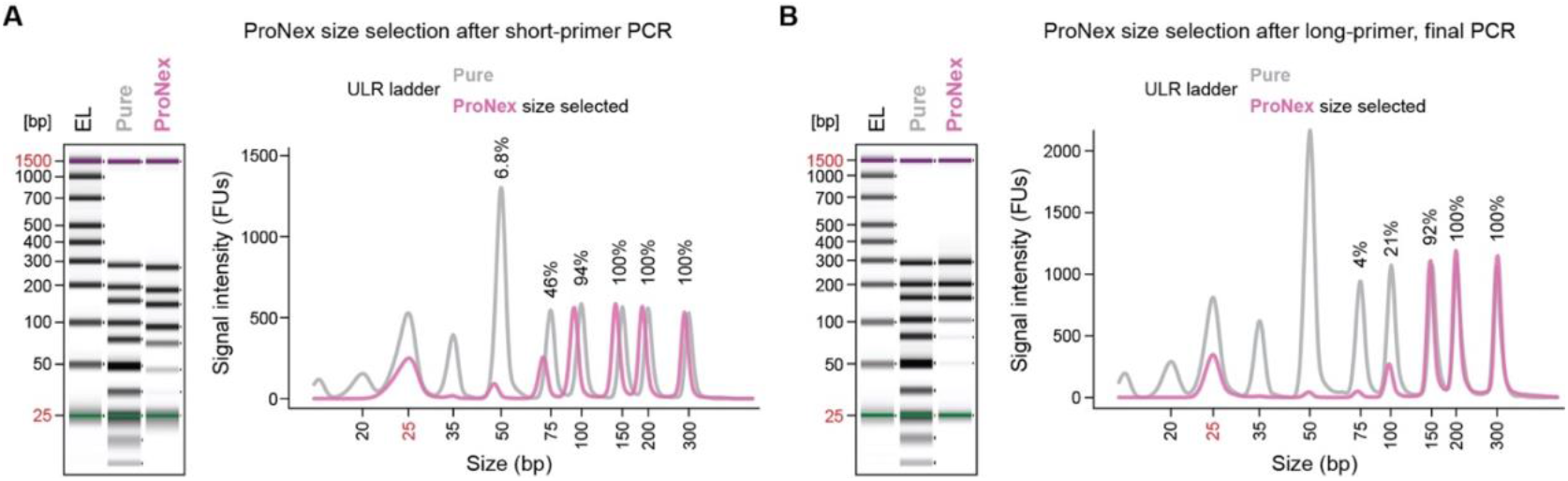
iCLIP3 library size selection. **(A)** ProNex bead-based size selection of dsDNA after short-primer PCR. Capillary gel electrophoresis image of the pure (grey) and size-selected ULR ladder (pink) using ProNex beads at sample-to-ProNex bead ratio of 1:2.95. **(B)** ProNex bead-based size selection of dsDNA after final long-primer PCR. Capillary gel electrophoresis image of the pure (grey) and size-selected ULR ladder (pink) using ProNex beads, PEG8000/NaCl mixture and ethanol. For both (A) and (B), accompanying partial electrograms of the capillary gel electrophoresis are shown. The numbers above peaks represent the percent recovery of each peak after ProNex bead size selection.

#### Long-primer test PCR

**Day 4 Timing: 30 min**

In this step, the short iCLIP3 library is amplified using long PCR primers containing the index sequences *i5* and *i7*, which are required for binding to the Illumina flow cell and for cluster generation during sequencing. Only an aliquot of the short iCLIP3 library is used in this test PCR, allowing both qualitative and quantitative assessment of the resulting test iCLIP3 library.

41. Long-primer test PCR.
  a. Assemble the following PCR reaction:

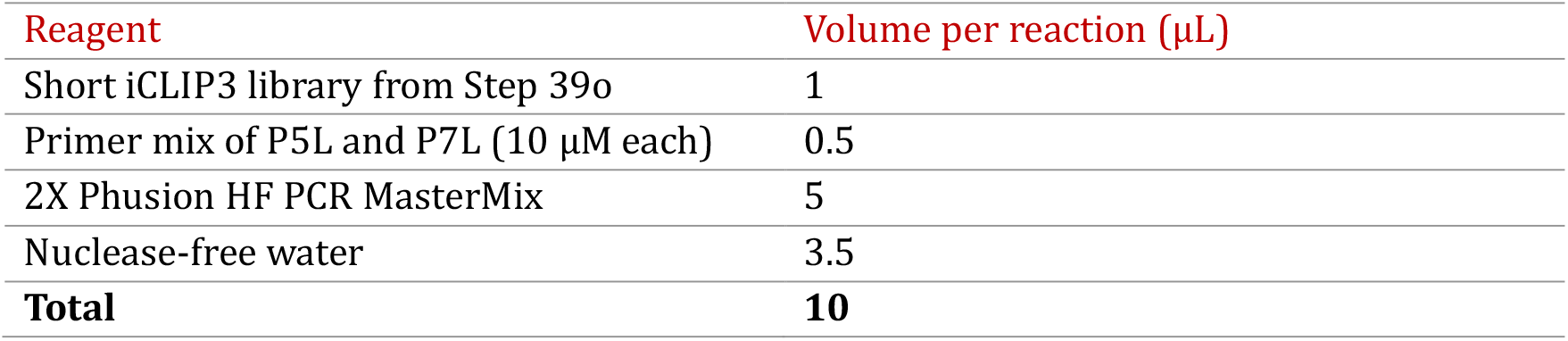
  b. Run the PCR in a thermocycler using the following PCR program:

b. Run the PCR in a thermocycler using the following PCR program:
| Steps | Temperature | Time | Cycle number |
| --- | --- | --- | --- |
| Denaturation 1 | 98°C | 30 s | 1 |
| Denaturation 2 | 98°C | 10 s | 8-15 |
| Annealing | 65°C | 30 s |  |
| Extension | 72°C | 30 s |  |
| Final extension | 72°C | 3 min | 1 |
| Hold | 16 | ∞ | 1 |

**Note:** Perform 2–3 PCR reactions for each sample using different cycle numbers (e.g., 8, 10, and 12 cycles) to determine the optimal amplification conditions for each library.

#### Test iCLIP3 library size selection with ProNex beads

**Day 4 Timing: 30 min**

In this step, ProNex beads are used to size-select the test iCLIP3 library. This step removes unused long PCR primers and recovers test iCLIP3 library fragments ≥ 170 bp (insert size: ≥ 25 bp).

42. Equilibrate ProNex beads and buffer (with ethanol added) at room temperature for 30 min prior to use.
43. Test iCLIP3 library size selection.
  a. Vortex ProNex bead stock and mix thoroughly by pipetting.
  b. Add 2.4 volumes of ProNex beads (24 µL) to the PCR reaction.
  c. Mix well by pipetting up and down 8-10 times.
  d. Incubate the mixture at room temperature for 10 min.
  e. Place the tubes on a magnetic rack to collect the beads.
  f. Discard the supernatants.
  g. Add 120 µL ProNex Wash buffer without disturbing the beads and incubate 15 s.
  h. Repeat steps f and g.
  i. Discard the supernatant and air-dry the beads until cracking becomes apparent. **Note**: Do not over-dry the beads.
  j. Remove the tubes from the magnetic rack and resuspend the beads in 12 µL nuclease-free water.
  k. Incubate the samples for 5 min at room temperature.
  l. Place the tubes on a magnetic rack to collect the beads.
  m. Transfer the eluted test iCLIP3 libraries to clean 1.5 mL tubes.

**Pause point**: The isolated test iCLIP3 libraries can be stored at 4ºC overnight or at -20ºC long term.

#### Test iCLIP3 library quality control and quantification

**Day 4 Timing: 30 min**

This step evaluates the quality and purity of the test iCLIP3 library and determines its concentration.

44. Test iCLIP3 library quality control using the TapeStation.
  a. Equilibrate Tape Station reagents at room temperature for 20–30 min prior to use.
  b. Load 2 µL purified test iCLIP3 library from Step 43m onto the TapeStation using High Sensitivity D1000 ScreenTape.
  c. Visualize library size distribution and confirm removal of PCR primers.
  d. Determine the average size of the library in bp. The purified library typically appears as a dsDNA smear ≥ 175 bp.
45. Quantification of test iCLIP3 library.
  a. Use 1–2 µL purified test iCLIP3 library to measure the DNA concentration (ng/µL) using a Qubit fluorimeter according to the manufacturer’s instructions.
  b. Calculate library molarity using the following equations:

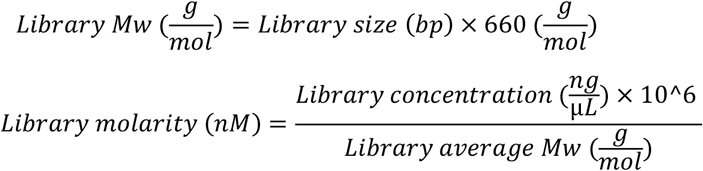

#### Long-primer final PCR cycle determination and primer selection

**Day 4 Timing: 30 min**

In this step, the optimal number of PCR cycles for final library amplification is determined. During the final PCR, each sample is labeled with unique dual indexes (UDIs) incorporated into the long primers. UDIs should be carefully selected to enable multiplexing with proper color balance and sufficient Hamming distance between indexes, thereby minimizing index hopping during sequencing.

46. The number of PCR cycles required for final library amplification can be calculated using the following equation:

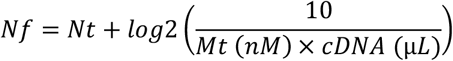

where *Nf* is the number of cycles for the final PCR, *Nt* is the number of cycles used in the test PCR, *Mt* is the calculated molarity from the test PCR, and cDNA is the volume of the pre-amplified cDNA (short dsDNA libraries) in the final PCR reaction.
47. For final PCR amplification of individual iCLIP3 libraries, unique dual index (UDI) primer pairs are selected with the DNA Barcode Combination Finder Shiny app (https://www.biozentrum.uni-wuerzburg.de/brb/resources/), which identifies compatible barcode combinations and ensures appropriate index color balance for Illumina sequencing. **Note:** The app takes as input the available *i7* (and optionally *i5*) index sequences, either from the built-in iCLIP3 index library or from user-provided custom index sets. It then calls the ‘experimentdesign’ function from the R package DNABarcodeCompatibility (Tre beau et al., 2019) to compute sets of barcode combinations that maintain a minimal pairwise distance (Hamming or Levenshtein) between indexes and thus reduce the risk of index misassignment or hopping. In addition, for Illumina platforms using XLEAP SBS chemistry (NextSeq 1000/2000 and NovaSeq X/X Plus), the app evaluates base composition per index cycle and iteratively searches for combinations that satisfy Illumina’s color-balance constraints (avoiding cycles with only A/G or only G signal, while allowing T/C-only cycles).
  a. Specify whether libraries will be single-indexed (*i7* only) or dual-indexed (*i7*+*i5*), and choose whether to use the iCLIP3 index library or paste custom index sequences (one per line).
  b. Enter the total number of libraries (samples), the desired multiplexing level per lane, and the number of color channels corresponding to the sequencing platform (e.g., 2 for NextSeq/NovaSeq).
  c. Optionally set advanced parameters by choosing the type and value for the distance metric (Hamming or sequence Levenshtein), which controls how dissimilar individual indexes must be.
  d. Run the analysis to obtain a table of optimal *i7* only or *i7*/*i5* UDI combinations, grouped by lane, along with summary tables showing per-cycle Index color balancing and sequence logos visualizing nucleotide-specific color signal and base usage across indexes.

The resulting table directly lists the recommended P5/P7 long primer pairs (containing *i5*/*i7* indexes) that should be used for the final PCR amplification of each iCLIP3 library.

#### Long-primer final PCR

**Day 4 or 5 Timing: 30 min**

In this step, the final iCLIP3 library is amplified using the primer combinations selected in the previous step. The resulting library is ready for sequencing on Illumina platforms.

48. Long-primer, final PCR.
  d. Assemble the following PCR reaction:

  d. Run the PCR in a thermocycler using the following PCR program:

| Reagent | Volume per reaction (μL) |
| --- | --- |
| Short iCLIP3 library from Step 39o | 10 |
| Primer mix of P5L and P7L (10 μM each) | 2 |
| 2X Phusion HF PCR MasterMix | 20 |
| Nuclease-free water | 8 |
| **Total** | **40** |

**Note:** Maintain the PCR reaction volume exactly at 40 μL.

- d. Run the PCR in a thermocycler using the following PCR program:
| Steps | Temperature | Time | Cycle number |
| --- | --- | --- | --- |
| Denaturation 1 | 98°C | 30 s | 1 |
| Denaturation 2 | 98°C | 10 s | n |
| Annealing | 65°C | 30 s |  |
| Extension | 72°C | 30 s |  |
| Final extension | 72°C | 3 min | 1 |
| Hold | 16 | ∞ | 1 |

#### Final iCLIP3 library size selection with ProNex beads

**Day 4 or 5 Timing: 30 min**

In this step, the final iCLIP3 library is purified using ProNex beads, and unused long PCR primers are removed. Low Range (ULR) DNA ladder is used to evaluate ProNex-based size selection.

49. Equilibrate ProNex beads and buffers, as well as PEG/NaCl Mix at room temperature for 30 min.
50. Final iCLIP3 library size selection.
  a. Set up the size selection ULR ladder control.

  b. Set up the reference ULR ladder and keep at 4ºC until further use:

| Reagent | Volume per reaction (μL) |
| --- | --- |
| Nuclease-free water | 19 |
| ULR DNA ladder | 1 |
| 2X Phusion HF MasterMix | 20 |
| **Total** | **40** |

  c. Prepare the following size selection mixture and pipette thoroughly to ensure proper mixing:

- b. Set up the reference ULR ladder and keep at 4°C until further use:
| Reagent | Volume per reaction (μL) |
| --- | --- |
| ULR DNA ladder | 1 |
| Nuclease-free water | 11 |
| **Total** | **12** |

  d. Add 40 µL size selection mixture to the PCR sample and to the size selection ULR ladder control.
  e. Add 20 µL 100% ethanol to the samples and mix by pipetting up and down 8-10 times.
  f. Incubate the mixture for 10 min at room temperature.
  g. Place the tubes on a magnetic rack to collect the beads.
  h. Discard the supernatant.
  i. Add 150 µL ProNex Wash buffer without disturbing the beads and incubate for 15 s.
  j. Repeat steps h and i.
  k. Discard the supernatant and air-dry the beads until cracking becomes apparent. **Note**: Do not over-dry the beads.
  l. Remove the tubes from the magnetic rack and resuspend the beads in 12 µL nuclease-free water.
  m. Incubate the tubes for 5 min at room temperature.
  n. Place the tubes on the magnetic rack to collect the beads.
  o. Transfer the eluted final iCLIP3 library and size selected ULR ladder control to clean 1.5 mL tubes.

- c. Prepare the following size selection mixture and pipette thoroughly to ensure proper mixing:
| Reagent | Volume per reaction (μL) |
| --- | --- |
| Nuclease-free water | 2.5 |
| PEG/NaCl Mix | 12.5 |
| ProNex beads | 25 |
| **Total** | **40** |

**Pause point**: The isolated final iCLIP3 library can be stored at 4ºC overnight or at -20ºC long term.

#### Evaluation of ProNex size selection for ULR ladder

**Day 4 or 5 Timing: 20 min**

This step evaluates the efficiency of the ProNex size selection using the reference ULR ladder and the size-selected ULR ladder control.

51. Evaluation of ProNex size selection using the TapeStation.
  a. Equilibrate TapeStation reagents at room temperature for 20–30 min prior to use.
  b. Load 2 µL reference ULR ladder from Step 50b and 2 µl size-selected ULR ladder control from Step 50o onto the TapeStation using High Sensitivity D1000 ScreenTape.
  c. Assess the recovery of the 75-bp, 100-bp and 150 bp-bands in the size-selected ULR ladder control relative to the reference ULR ladder.

**Note**: Successful size selection is indicated by significant removal of the 75-bp band and ≥ 90% recovery of the 150-bp band (**Fig. 4B**). Additionally, the ratio of the 150-bp to 75-bp band intensities in the size-selected ULR ladder control should be ≥ 15.

#### Final iCLIP3 library quality control and quantification

**Day 4 or 5 Timing: 30 min**

This step evaluates the quality, purity and concentration of the final iCLIP3 library.

52. Final iCLIP3 library quality control using the TapeStation.
  a. Equilibrate TapeStation reagents at room temperature for 20–30 min prior to use.
  b. Load 2 µL final iCLIP3 library from Step 50o on the TapeStation using High Sensitivity D1000 ScreenTape.
  c. Visualize the library size distribution and confirm the removal of PCR primers.
  d. Determine the average library size for each sample in bp. A properly purified library appears as a dsDNA smear ≥ 175 bp.
  e. Quantify the final iCLIP3 library using Qubit fluorimeter or equivalent instrument.
53. Quantification of final iCLIP3 library.
  c. Use 1 µL final iCLIP3 library to measure the concentration (ng/µL) for each sample with a Qubit fluorimeter according to the manufacturer’s instructions.
  c. Using the equations from Step 45b, determine the molarity of each individual iCLIP3 library.

**Pause point**: The isolated final iCLIP3 libraries can be stored at -20ºC long term.

#### iCLIP3 library multiplexing

**Day 4 or 5 Timing: 20 min**

In this step, multiple individual iCLIP3 libraries are combined at equimolar ratio to enable sequencing on the same Illumina flow cell lane. Equimolar pooling ensures that each library is represented at comparable sequencing depth within the pooled sample.

54. Using the library molarity calculated in Step 53b, mix the individual libraries at equimolar ratios to create the final multiplexed library pool.

#### Pooled iCLIP3 library cleanup with ProNex beads

**Day 4 or 5 Timing: 30 min**

This step removes residual long PCR primers from the pooled iCLIP3 library, ensuring a clean library for sequencing.

55. Purify the pooled iCLIP3 library by following Steps 42–45 of this protocol.

**Pause point**: The purified pooled iCLIP3 library can be stored at -20ºC long term until sequencing.

#### Pooled iCLIP3 library quality control and quantification

**Day 4 or 5 Timing: 30 min**

This step assesses the quality, size distribution and concentration of the pooled iCLIP3 library to ensure it is suitable for sequencing.

56. Evaluate the pooled iCLIP3 library using the TapeStation to visualize the size distribution and confirm the removal of residual primers (follow Step 52). Measure the concentration of the pooled iCLIP3 library using Qubit fluorimeter or an equivalent instrument (follow Step 53).

### EXPECTED OUTCOMES

Using Protocol 2, iCLIP3 libraries are expected to be ≥ 175 bp, with a majority of the signal falling within a size range of **175–300 nt**. No signal should be detectable in negative control samples, including UV– and no-antibody controls, indicating minimal background and high specificity of the procedure. Experiments in which the long-primer final PCR exceeds 15 cycles are expected to yield substantial levels of PCR duplicates and may therefore not be suitable for downstream analyses.

Using this protocol, iCLIP3 libraries were successfully prepared for the RNA-binding protein U2AF2 from HeLa cell lysates containing 250 µg, 100 µg, and 40 µg of total protein. Long-primer final PCR amplification with 5, 6, and 7 cycles yielded high-quality libraries suitable for sequencing (**Fig. 5A, B**).

**Figure 5.**
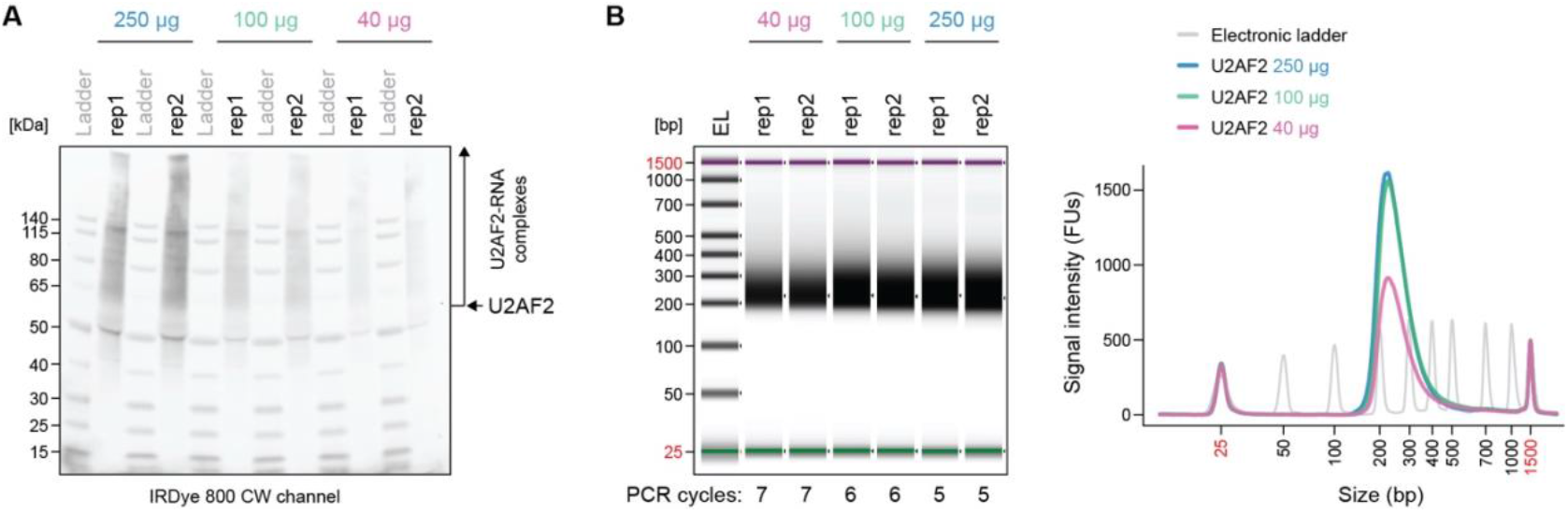
Generation of U2AF2 iCLIP3 libraries from different amounts of input material. **(A)** Nitrocellulose membrane showing U2AF2–RNA complexes immunopurified from HeLa cell lysates containing 250 µg, 100 µg and 40 µg total protein. Only 10% of the immunopurified U2AF2–RNA complexes in each sample and each replicate were labeled with pCp-IR750 and visualized in the IRDye 800 CW channel. The remaining 90% of the U2AF2–RNA complexes were subjected to L7 linker ligation. Both 10% and 90% fractions were combined and ran in the same gel lane. **(B)** Left panel depicts TapeStation capillary gel electrophoresis with the final iCLIP3 libraries prepared from the samples depicted in (A). The number of cycles used in the second, final PCR is shown. Right panel shows an accompanying electrogram of the capillary gel electrophoresis from the left panel. The average signal intensity values from two replicates are displayed. FUs, fluorescent units.

### LIMITATIONS

Like other CLIP-based protocols, iCLIP3 depends on the availability of high-quality IP-grade antibodies, which can limit its application to proteins for which suitable antibodies are lacking. In such cases, epitope tagging of the protein of interest can help overcome this limitation. iCLIP3 is also constrained by the inherently low efficiency of UV irradiation to induce covalent crosslinking between proteins and RNA; consequently, some RBPs may fail to crosslink efficiently despite bona fide interactions with RNA. For these proteins, metabolic labeling with 4-thiouridine (4sU) followed by 365-nm UV crosslinking may improve crosslinking efficiency, but different computational analysis of the data would have to be applied (Hafner et al., 2010). Importantly, the iCLIP3 protocol cannot identify binding sites of cellular proteins whose interactions with RNA do not involve direct protein–RNA contact. In such cases, alternative approaches such as RIP-seq can provide complementary insights into RNA–protein associations.

### TROUBLESHOOTING

#### Problem 1

No PCR libraries are observed in the test PCR with 12 cycles.

#### Potential solution

Increase the number of PCR cycles. The iCLIP3 library is likely derived from negligible amounts of immunopurified RNA. Although additional PCR cycles may produce a visible library with sufficient yield, a high rate of PCR duplication is expected. For optimal results, repeat the experiment using a substantially larger amount of starting material, such as a higher number of cells.

#### Problem 2

The peak observed in the test PCR is around **150 bp**, indicating the absence of cDNA inserts or the presence of only very short inserts.

#### Potential Solution

This issue is similar to that described in Problem 1 and likely reflects a failure during library preparation. Possible causes include insufficient input material. Additional factors may include suboptimal reaction conditions, overdigestion of the samples, or excessive RNase activity. Excessive RNase activity could lead to RNA degradation and consequently very short or absent cDNA inserts. To improve outcomes, the experiment should be repeated using a substantially larger amount of starting material, such as a higher number of cells. Reducing the amount of RNase or shortening the RNase incubation time may also help preserve RNA integrity. If the problem persists after repetition, the reagents should be replaced. To facilitate this, all reagents should be aliquoted to avoid repeated freeze–thaw cycles.

#### Problem 3

Additional bands at higher molecular weight are visible in the library profiles.

#### Potential Solution

This pattern likely indicates overamplification and the formation of secondary PCR products. To reduce the occurrence of these higher-molecular-weight bands, the number of cycles in the long-primer final PCR should be reduced.

#### Problem 4

Signal is detected in the UV– and no-antibody control samples.

#### Potential solution

This signal may arise from non-specific background. Increasing the stringency and number of wash steps may help reduce background, and protein–RNA complexes should be carefully monitored on the membrane in the UV– and no-antibody control lanes. Alternatively, the signal may result from contamination with material from previous experiments. In this case, thorough cleaning of the workspace is recommended, along with strict physical separation of pre- and post-PCR work areas to minimize cross-contamination.

## PROTOCOL 3 PROCESSING OF iCLIP3 SEQUENCING DATA WITH RACOON_CLIP

After high-throughput sequencing of the iCLIP3 libraries, the resulting data are used to derive transcriptome-wide RNA–protein crosslink profiles. Since the sequencing reads follow a particular design and harbor protocol-specific elements such as UMIs, dedicated preprocessing strategies are needed (Chakrabarti et al., 2018). To achieve this, we employ racoon_clip (Klostermann and Zarnack, 2024), a fully automated workflow for processing iCLIP3, earlier iCLIP variants and eCLIP data to extract RNA–protein crosslink positions at single-nucleotide resolution. We have previously applied racoon_clip, for example, to identify microRNA-dependent AGO binding sites in mouse cells (Verheyden et al., 2024).

In brief, the pipeline begins with preprocessing steps, include optional quality filtering and trimming of UMIs and sequencing adapters, that are tailored to different CLIP protocols to ensure correct parsing of read structures. Processed reads are then aligned to the reference genome in a splice-aware manner using STAR (Dobin et al., 2013) with parameters optimized to preserve exact 5′ ends, followed by UMI-based deduplication to remove PCR duplicates. Crosslink sites are subsequently defined as the nucleotide immediately upstream of aligned read 5′ ends and exported as single-nucleotide resolution tracks in BIGWIG format. In the latest version used here, the workflow is extended by automated peak calling with PureCLIP to identify crosslink positions significantly enriched over local background; these peaks form the basis for the downstream binding site definition (see Protocol 4 below). In addition, racoon_clip generates a comprehensive HTML report summarizing data quality metrics and processing statistics.

### STEP-BY-STEP METHOD DETAILS

#### Installation of racoon_clip

**Timing: 1-2 h**

1. Install racoon_clip on a UNIX-based compute cluster.
  a. Build racoon_clip in a conda environment from the GitHub release. **Note:** racoon_clip is also available as ready-made containers based on Docker or Apptainer. For more details, refer to the racoon_clip documentation (https://racoon-clip.readthedocs.io/en/latest/methods_description.html). SingularityCE is currently not supported.
    i. Make a fresh conda environment containing the racoon_clip dependencies mamba, python and pip.

~~~
<u>bash</u>
# make racoon_clip conda environment with mamba support
> conda create -n racoon_clip \
   --override-channels -c conda-forge \
   mamba=1 \
   ‘python_abi=*=*cp*’ \
   python=3.9.0 \
   pip=25.0
# activate conda environment
>conda activate racoon_clip
~~~
    ii. Find the latest version at https://github.com/ZarnackGroup/racoon_clip/releases and copy the link to the ZIP file of the lates version.
    iii. Download and install racoon_clip in this environment. **Note:** Replace [version] with the latest version number (e.g., v.2.0.11).

~~~
<u>bash</u>
# download racoon_clip
>wget link/to/latest/release.zip
>unzip [version].zip
>cd racoon_clip-[version]
# install
pip install -e .
~~~
  c. Check that racoon_clip works by getting the version number. In this protocol, we are using version v.2.0.12.

~~~
<u>bash</u>
>racoon_clip -v
~~~ **Optional:** Test racoon_clip. This runs a quick test including basic functionality checks and should output the message “All tests passed successfully” in the end. If not, see the troubleshooting section below.

~~~
<u>bash</u>
>racoon_clip test --light
~~~

#### Preparation of genome input files

**Timing: 1-2 h**

2. Download a suitable genome (FASTA format) and a matching genome annotation in GTF format. For human or mouse data, we recommend using the comprehensive gene annotation for the primary genome assembly (PRI) from GENCODE (https://www.gencodegenes.org).

~~~
<u>bash</u>
# download genome FASTA and annotation GTF
>wget link_to_genome.fa.gz
>wget link_to_genome_annotation.gtf.gz
~~~ **Optional:** If you want to use racoon_clip’s FastQ screen module to screen for potential RNA contamination from other organisms and/or rRNA content, you need to download these genomes or rRNA and prepare mapping indices of them.
  a. Contaminant organisms: Get FASTA files of all genomes you want to screen against.

~~~
<u>bash</u>
# download genomes to screen against
>wget link_to_genome_for_screening.fa.gz
>fa=<path/to/genome_for_screening.fa.gz>
~~~
  b. rRNA: Get a FASTA file from the rRNA sequences. A FASTA file of the human rRNA sequences used for this study can be found on GitHub (https://github.com/ZarnackGroup/Despic_et_al_2026). Download it there or make your own by following steps:
    i. Make a new empty FASTA file (e.g., rRNA.fa).
    ii. Copy the rRNA sequences for example from NCBI (https://www.ncbi.nlm.nih.gov/gene/) into the FASTA file. Follow the FASTA format as shown here.

~~~
<u>**content of rRNA.fa**</u>
>28S_rRNA
CGCGACCTCAGATCAGACGTGGCGACCCGCTGAATTTAAGCATA…
>18S_rRNA
…
~~~ **Note:** For the human genome, we used the rRNA genes from NCBI (https://www.ncbi.nlm.nih.gov/gene/) with the following NCBI/RefSeq IDs: 28S, NR_003287.4; 18S, NR_003286.4; 5.8S, NR_003285.3; 5S, NR_023363.1; mitochondrial 12S (MT-RNR1), GeneID 4549; mitochondrial 16S (MT-RNR2), GeneID 4550.
  c. Generate genome indices for mapping.
    i. Install necessary packages in a conda environment.

~~~
<u>bash</u>
# install bowtie and gffread
>conda create -n bowtie_index bioconda::bowtie2
>bioconda::gffread
>conda activate bowtie_index
>bowtie2 -h
>gffread -h
~~~
    ii. Index the genomes with bowtie2.
  d. If you want to screen for multiple contaminants and/or the rRNAs of multiple organisms, repeat the steps a/b and c for each of them. Create a configuration file named fastqscreen.config with all contaminants to screen against. Use the following space-separated structure (DATABASE <name_of_contaminant> <path_to_folder_containing_bowtie_index/prefix_of_bowtie_index>), adding a new line with the same structure for each contaminant to be screened:

~~~
<u>**content of fastqscreen.config:**</u>
DATABASE contaminantA </path/to/contaminantA_idx>
DATABASE rRNA </path/to/rRNA_idx>
…
~~~

#### Obtaining crosslink events and peaks from sequencing data using racoon_clip

**Timing: 2-3 days**

In this step, the raw sequencing data is processed with the command-line tool racoon_clip. In brief, racoon_clip performs quality checks, adapter trimming, genomic alignment, deduplication and extraction of both crosslink events and peaks at single-nucleotide resolution (**Fig. 6A**). The default parameters for all steps can be found in the racoon_clip documentation (https://racoon-clip.readthedocs.io/en/latest/methods_description.html).

**Figure 6.**
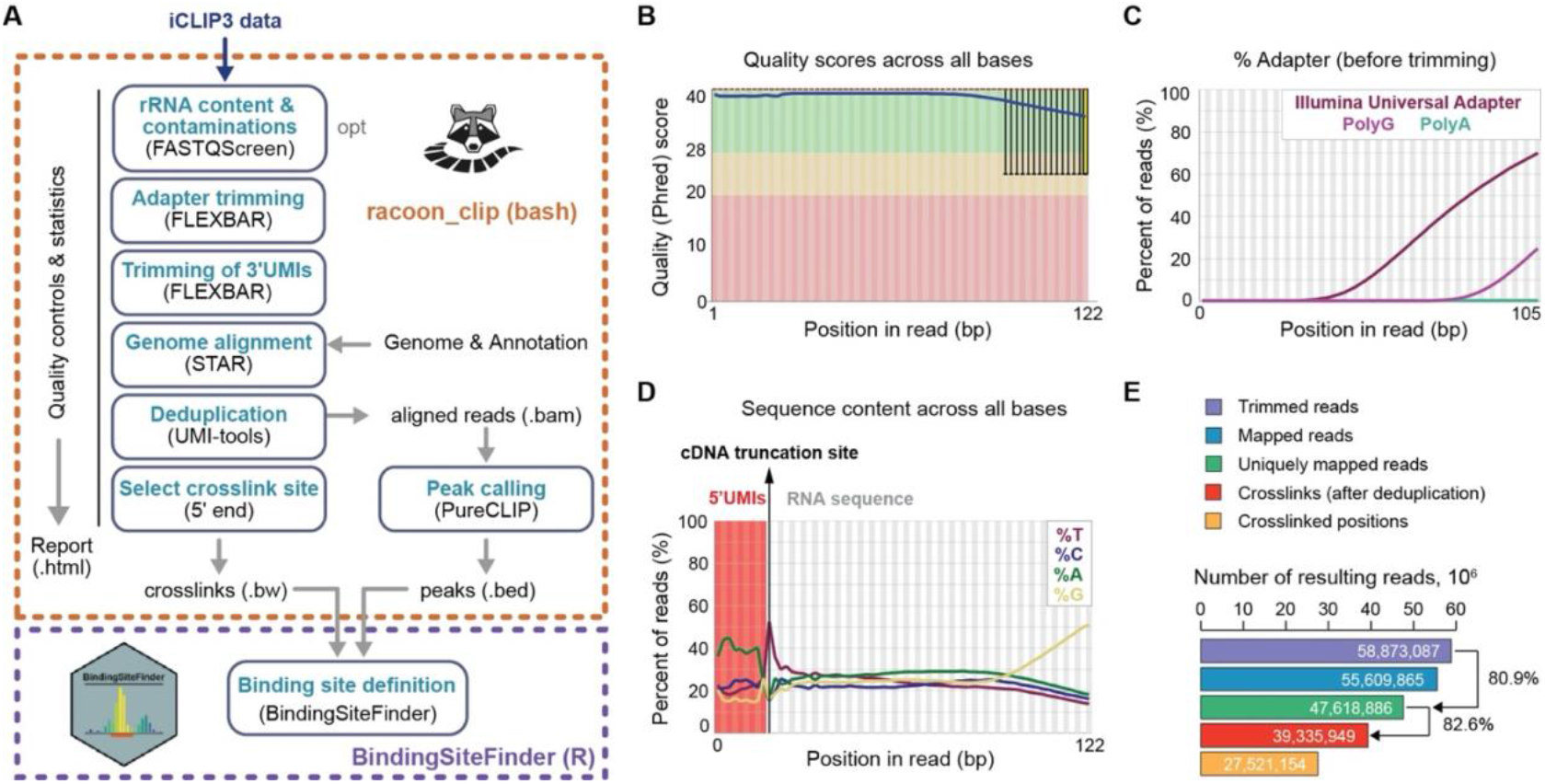
Processing of iCLIP3 sequencing data. **(A)** Computational analysis workflow for iCLIP3 data using the command-line tool racoon_clip and the R/Bioconductor package BindingSiteFinder, including quality control, identification of crosslink sites, peaks and binding sites. Shown are major steps with tools used (in brackets) and output files. **(B)** Distribution of sequencing quality (Phred) scores at each nucleotide position along iCLIP3 reads. **(C)** Adapter content before trimming. Plot shows percent of reads with universal Illumina adapter, polyG or polyA tracts at each nucleotide position in reads. **(D)** Sequence composition of reads. Shown is the percent of each nucleotide at each nucleotide position. Position of 9-nt 5′UMI and cDNA truncation site are highlighted. **(E)** Number of reads after each racoon_clip processing step. (B–E) show exemplary visualizations (modified from racoon_clip report) for replicate 1 of 250 µg U2AF2 iCLIP3. The full processing report for all U2AF2 iCLIP3 samples is available in Supplementary Material 1.

**Note:** Scripts and configuration files used for the U2AF2 iCLIP3 data in this publication are available on GitHub (https://github.com/ZarnackGroup/Despic_et_al_2026).

3. Prepare the racoon_clip input files.
  a. Copy the raw demultiplexed FASTQ sequencing files to the compute cluster. **Note:** racoon_clip can handle gzipped or unzipped FASTQ files. We recommend comparing the md5sums to ensure that the transfer was complete. **Optional:** Make an adapter file (here named adapter.fa) that contains the sequence of the L7 adapter:

~~~
<u>**content of adapter.fa**</u>
>adapter
AGATCGGAAGAGCACACGTC
~~~ **Note:** If you are using an adapter that differs from the one used in this protocol, add the sequence of your adapter instead.
  b. Make a racoon_clip config file (here called config.yaml) in YAML format that specifies all input files and selected parameters.
    i. A basic racoon_clip config file looks like this:

~~~
<u>**content of racoon_config.yaml**</u>
# output directory
wdir: “output/path”
# input
infiles: “path/to/sample1.fastq path/to/sample2.fastq”
samples: “sample1 sample2”
# annotation
gtf: “path/to/annotation.gtf”
genome_fasta: “path/to/genome_assembly.fa”
read_length: N
# experiment type
experiment_type: “iCLIP3”
# adapter
adapter_trimming: True
adapter_file: “path_to/adapter.fa”
~~~
    ii. Fill the config file as follows:

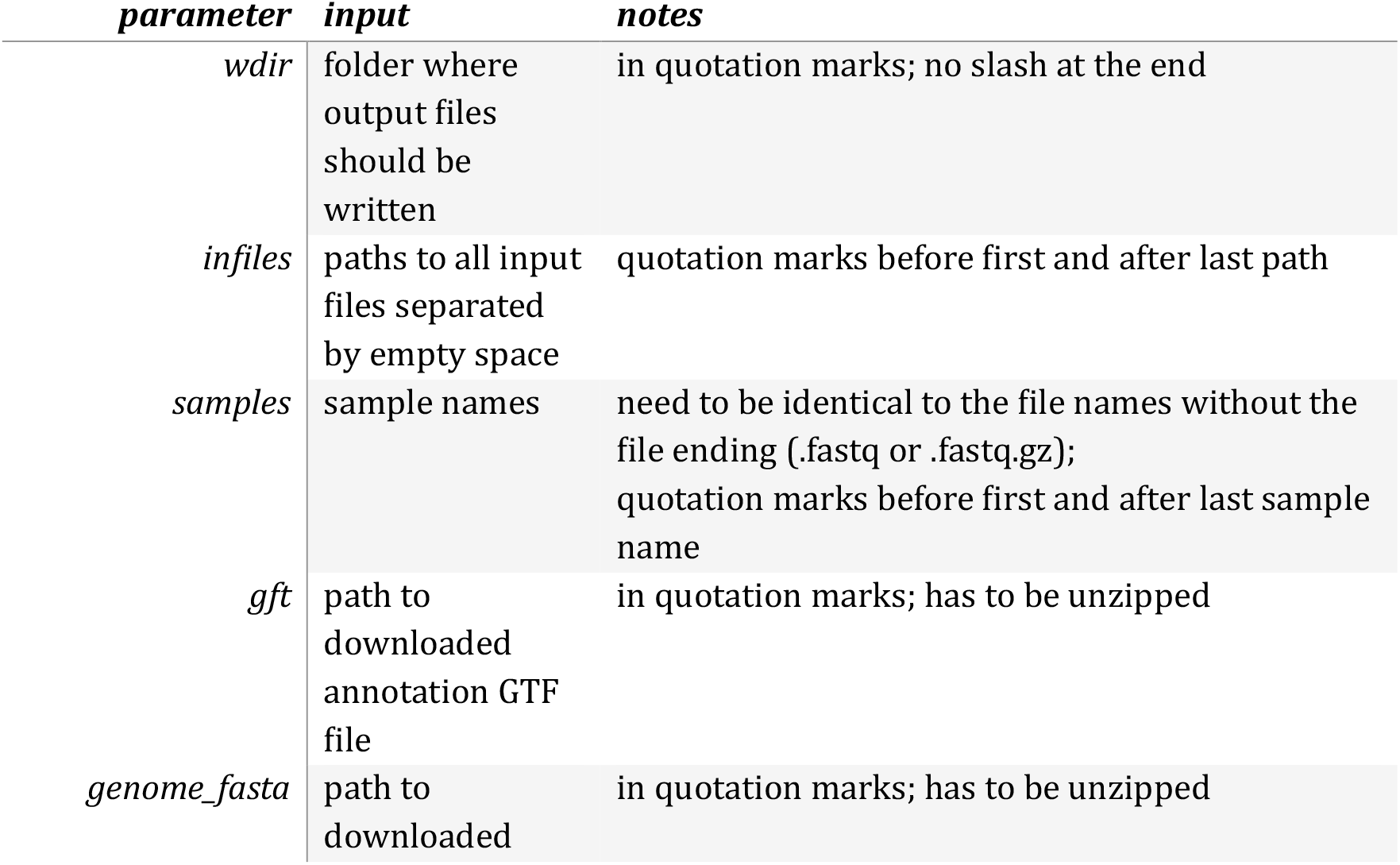
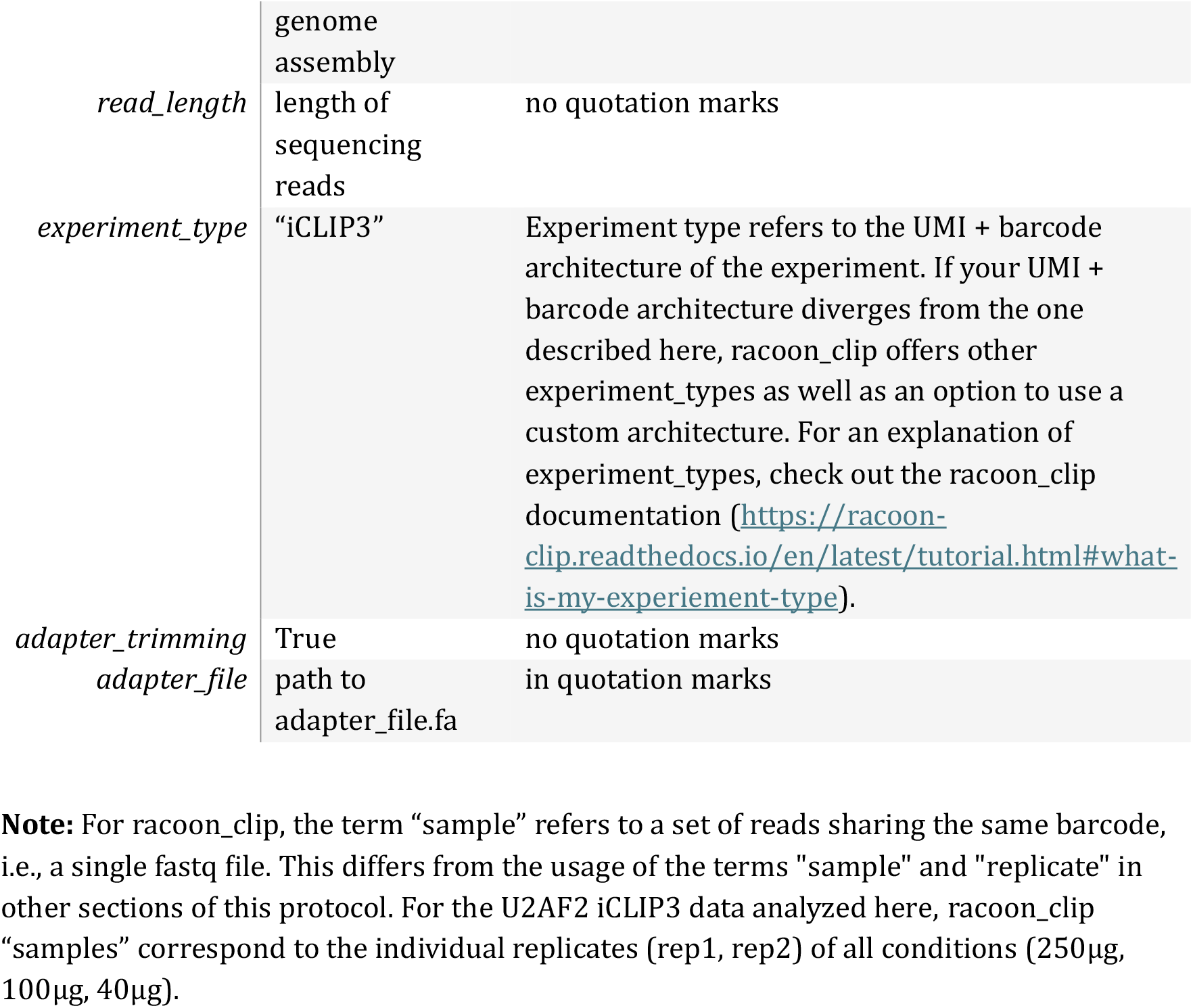 **Note:** For racoon_clip, the term “sample” refers to a set of reads sharing the same barcode, i.e., a single fastq file. This differs from the usage of the terms “sample” and “replicate” in other sections of this protocol. For the U2AF2 iCLIP3 data analyzed here, racoon_clip “samples” correspond to the individual replicates (rep1, rep2) of all conditions (250µg, 100µg, 40µg).
  c. racoon_clip can be customized by adding additional parameters as extra lines to the config file. The following lists the commonly used parameters. A complete description can be found in the racoon_clip documentation (https://racoon-clip.readthedocs.io/en/latest/all_options.html). Customization 1 – Experiment groups: If your data contains multiple conditions or groups, you can perform peak calling separately within each group.
    i. Prepare a group TXT file (here named groups.txt) to assign each sample to a group. The group file should have one line per sample and specify “group_name sample” in each line (separated by a space). **Important:** The sample names need to be exactly the file names without file ending (.fastq or .fastq.gz). Group names can be chosen freely. Here is an example for the U2AF2 iCLIP3 data from this study:

~~~
<u>**content of groups.txt**</u>
u2af2_40ug u2af2_40ug_rep1.R1
u2af2_40ug u2af2_40ug_rep2.R1
u2af2_100ug u2af2_100ug_rep1.R1
u2af2_100ug u2af2_100ug_rep2.R1
u2af2_250ug u2af2_250ug_rep1.R1
u2af2_250ug u2af2_250ug_rep2.R1
~~~ **Note:** If you do not specify a group file, peak calling will be performed on a merge of all samples.
    ii. Add the following to the config.yaml.

~~~
<u>**optional lines for racoon_config.yaml**</u>
experiment_group_file: “path/to/groupfile.txt”
~~~ Customization 2 – Contamination screening: If you want to screen for potential contaminations, set the parameter fastqScreen to True and provide the path to the fastqscreen.config file to the parameter fastqScreen_config.

~~~
<u>**optional lines for racoon_config.yaml**</u>
fastqScreen: True
fastqScreen_config: “path_to/fastqscreen.config
~~~ Customization 3 – Restrict PureCLIP training on selected chromosomes (Recommended): Peak calling with PureCLIP (Krakau et al., 2017) is the most time and memory-consuming step of racoon_clip. PureCLIP first trains a hidden markow model on the data and then calls peaks using the learned pattern. Training is by default done on the complete genome, but can be downsized to a few chromosomes, as also recommended in the PureCLIP documentation (https://pureclip.readthedocs.io/en/latest/PureCLIPTutorial/basicMode.html). To restrict PureCLIP training to the first 3 chromosomes, add the following line including a semicolon-separated list of the chromosome names to the racoon_clip config file.

~~~
<u>**optional lines for racoon_config.yaml**</u>
morePureclipParameters: “-iv ‘chr1;chr2;chr3;’“
~~~ **Note:** Chromosome naming might differ depending on the genome and source (e.g., ‘1;2;3;’ is often used by ENSEMBL). The results shown in **Fig. 6–8** were obtained by training on the full genome.
4. Run racoon_clip by specifying the path to the config file and the number of available CPUs.

~~~
<u>bash</u>
> racoon_clip peaks --configfile <your_configfile.yaml> --
cores <n_cores>
~~~ **Note:** This step will likely take > 36 h. Increasing <n_cores> can speed this up to a certain extent. As a rule of thumb, it makes sense to specify the number of CPUs as 1x, 2x or 4x the number of samples. For the U2AF2 iCLIP3 data, computation time for all samples was around 2 days with 20 CPUs. **Optional:** If your compute cluster is based on a SLURM architecture, you can provide a cluster profile to start the racoon_clip processing steps as individual jobs. Instructions can be found in the racoon_clip documentation (https://racoon-clip.readthedocs.io/en/latest/all_options.html#cluster-execution).
5. racoon_clip will produce a folder called results inside the output directory specified in the config file. Download the following files/folders to a local computer to continue with the protocol below:
  a. the file Report.hmtl, which summarizes the processing steps and quality controls performed by racoon_clip,
  b. the folder bw, which contains the crosslink events of each sample in BIGWIG format,
  c. the folder bw_merged, which contains the merged crosslink events for each group (or across all samples if no group was specified) in BIGWIG format,
  d. the folder peaks, which contains BED files with the PureCLIP peaks for each group (or one file for the merge of all samples if no group was specified).
6. Check the Report.html file to control the quality of the data and processing along the performed steps.

### EXPECTED OUTCOMES

racoon_clip generates a full report (in HTML format) that documents the performance and data quality across the analysis workflow. Exemplary visualizations for sequencing quality, adapter content, sequence composition and genomic mapping results for one sample are shown in **Fig. 6B–E**. The full processing report for all U2AF2 iCLIP3 samples is available in **Supplementary Material 1**.

The main output of racoon_clip are the BIGWIG files containing the crosslink events for each sample as well as merged across all samples (or within each group if specified). The crosslink positions correspond to the nucleotide position 1 nt upstream of the reverse transcription termination sites. The crosslink tracks can be loaded into a genome browser (e.g., Integrative Genomics Viewer [IGV, https://igv.org/]) for manual inspection and comparison between replicates.

### LIMITATIONS

racoon_clip requires 120 GB of RAM during the genomic mapping step. Therefore, it cannot be run on most local computers, but requires access to a compute cluster, cloud computing or a special workstation.

A reference genome and annotation in standard GTF format are required for both the initial data processing with racoon_clip and the subsequent binding site definition with BindingSiteFinder (see below). These are not always available, particularly for non-model organisms.

### TROUBLESHOOTING

#### Problem 1

Low rate of uniquely mapping reads to the reference genome.

#### Potential solution

Poor unique mapping rates may result from experimental problems, such as RNA over-digestion, inefficient removal of adapter dimers or other contamination. Moreover, a high content of multimapping reads may originate from rRNA or repetitive sequences which are often contaminants but can also genuine RNA targets, depending on the RNA-binding protein of interest. Inspect the racoon_clip report to find out about potential problems and repeat the experiment if necessary.

#### Problem 2

High fraction of reads lost during duplicate removal.

#### Potential solution

A high duplication rate indicates a low library complexity which may originate from insufficient amounts of immunopurified RNA. This is usually reflected in an increased number of PCR cycles required during library preparation (see above).

#### Problem 3

The racoon_clip pipeline breaks during the peak calling step with PureCLIP.

#### Potential solution

This usually occurs if RAM limits are reached. Restrict the training step of PureCLIP to selected chromosomes (see above) and/or provide additional RAM.

#### Problem 4

racoon_clip fails at the start of the run.

#### Potential solution

This often points to either a problem with mamba or errors in the config file. Double check for typos in the file names and paths. If problems persist, post an issue on the GitHub page of racoon_clip (https://github.com/ZarnackGroup/racoon_clip/issues).

## PROTOCOL 4 DEFINING BINDING SITES WITH BINDINGSITEFINDER

### STEP-BY-STEP METHOD DETAILS

**Timing: 1-2 days**

After extracting the crosslink events with racoon_clip, the challenge is to define discrete binding sites among the broad background across transcripts. In most cases, the binding sites of the RNA-binding protein can be distinguished from background because they typically display higher signal and a bell-shaped accumulation of crosslink events (Sugimoto et al., 2012; Busch et al., 2020).

Within the racoon_clip workflow (Klostermann and Zarnack, 2024), enrichment of crosslink events over background (peak calling) is assessed using PureCLIP (Krakau et al., 2017). The resulting peak locations are provided as one BED file per sample and represent individual nucleotide positions with significantly enriched crosslink signal. These can be used as a basis for defining complete binding sites, which usually span 5–9 nt. In addition, robust binding site definition benefits from incorporating reproducibility across biological replicates.

We therefore recommend defining discrete, equally sized binding sites based on the racoon_clip output and applying a reproducibility filter. For this purpose, we use the R/Bioconductor package BindingSiteFinder (Bru ggemann et al., 2021), which refines binding site boundaries and supports replicate-aware filtering to improve robustness. The package provides a wrapper function that executes the complete workflow with automatic parameter estimation when no arguments are supplied. In addition, multiple parameters can be specified to refine binding site definition. We recommend testing different parameter settings and evaluating their impact by visualizing the resulting binding sites together with the racoon_clip crosslink tracks in a genome browser.

#### Setup of R and R packages

1. Install R software and the necessary R packages. **Note:** The definition of binding sites in R can be performed on a local computer.
  a. Install the latest version of R by following the installation guide: https://www.r-project.org/. **Optional:** Install the latest version of RStudio by following the installation guide: https://posit.co/download/rstudio-desktop/.
  b. Open RStudio or R and install the required R packages.

~~~
<u>R (all code blocks from here)</u>
if (!require(“BiocManager”, quietly = TRUE))
   install.packages(“BiocManager”)
install.packages(“knitr”)# optional for rendering reports
install.packages(“tidyverse”)
install.packages(“purrr”)
BiocManager::install(“rtracklayer”)
BiocManager::install(“BindingSiteFinder”)
BiocManager::install(“GenomicRanges”)
BiocManager::install(“GenomeInfoDb”)
BiocManager::install(“txdbmaker”)
~~~
2. Open a new file (e.g., in R, RMD or QMD format) to document the code for the analysis steps. **Note:** All following steps are provided in the example script in **Supplementary Material 2**.
3. Load the libraries.

~~~
# ----------------------
# load libraries
# ----------------------
library(knitr)
library(purrr)
library(rtracklayer)
library(tidyverse)
library(BindingSiteFinder)
library(GenomicRanges)
library(GenomeInfoDb)
library(GenomicFeatures)
library(txdbmaker)
~~~

#### Automatic binding site definition

4. Define an output folder.

~~~
# set output folder
out <- “<path/to/your/output_folder>“
~~~
5. Extract genes and gene regions from the annotation file as input for BindingSiteFinder. **Note:** The following steps need to be executed only once to extract the gene and transcript region coordinates and save them as RDS files. In all subsequent runs, annotations can be loaded directly from the RDS files to save compute time.
  a. Import annotation file (in GTF format). Ideally, the same annotation file as in racoon_clip should be used.

~~~
# ----------------------
# prepare annotation
# ----------------------
# GTF annotation file (downloaded for example from GENCODE)
annoFile <- “path/to/annotation.gtf.gz”
# make annotation database from GTF file
annoDb = txdbmaker::makeTxDbFromGFF(file = annoFile, format
= “gtf”)
annoInfo = rtracklayer::import(annoFile, format = “gtf”)
~~~
  b. Extract gene coordinates including metadata and save as RDS file.

~~~
# get genes as Granges
gns = genes(annoDb)
idx = match(gns$gene_id, annoInfo$gene_id)
meta = cbind(elementMetadata(gns),
             elementMetadata(annoInfo)[idx,])
meta = meta[,!duplicated(colnames(meta))]
elementMetadata(gns) = meta
out_gns <- paste0(out, “gns.rds”)
saveRDS(gns, out_gns)
~~~
  c. Extract transcript regions and save as RDS file.

~~~
# get transcript regions as Granges
cdseq = cds(annoDb)
intrns = unlist(intronsByTranscript(annoDb))
utrs3 = unlist(threeUTRsByTranscript(annoDb))
utrs5 = unlist(fiveUTRsByTranscript(annoDb))
regions = GRangesList(CDS = cdseq, INTRON = intrns, UTR3 =
utrs3, UTR5 = utrs5)
out_regions <- paste0(out, “regions.rds”)
saveRDS(regions, out_regions)
~~~
6. Run BindingSiteFinder in its default mode with automatic parameter estimation.
  a. Define the input files. Provide the path to the peak file (called by PureCLIP, in BED format) and to the folder with the crosslink files (in BIGWIG format). In the code example below, all samples in this folder will be used. **Note:** If you specified multiple groups in racoon_clip, binding sites should be defined separately for each group. This can be achieved by selecting only the files for the given condition or separating files into distinct folders.

~~~
# ----------------------
# get input from racoon_clip
# ----------------------
# PureCLIP file
pureclip_file <-
“<path/to/racoon_clip_out_folder/results/peaks/
pureclip_sites.bed>“
# crosslink files (if all samples are from the same
condition and you did not specify groups in racoon_clip)
bw_dir <- “path/to/racoon_clip_out_folder/results/bw”
bw.plus <- list.files(bw_dir, pattern = “plus.bw$”,
full.names = TRUE, recursive = TRUE)
bw.minus <- list.files(bw_dir, pattern = “minus.bw$”,
full.names = TRUE, recursive = TRUE)
~~~
  b. Load RDS files with stored gene and transcript regions from the annotation.

~~~
# read prepared annotation
gns <- readRDS(paste0(out, “gns.rds”))
regions <- readRDS(paste0(out, “regions.rds”))
~~~
  c. Import the peaks, clean up the peaks object and check the number of peaks. **Sanity check:** This number should be the same as given in the chapter “Peak calling” of the Report.html from racoon_clip.

~~~
# ----------------------
# peaks from PureCLIP
# ----------------------
# import PureCLIP peaks
peaks = rtracklayer::import(con = pureclip_file,
                             format = “BED”,
                             extraCols=c(“additionalScores”
= “character”))
# clean PureCLIP peaks columns
peaks$additionalScores = NULL
peaks$name = NULL
# check number of peaks
~~~ **Optional:** Remove scaffold chromosomes and the mitochondrial chromosome.

~~~
# optional: keep only standard chromosomes and drop chrM
peaks = keepStandardChromosomes(peaks, pruning.mode =
“coarse”) %>%
      dropSeqlevels(., “chrM”, pruning.mode = “coarse”)
~~~
  d. Generate a dataframe with the metadata for BindingSiteFinder. Check the printed metadata dataframe and make sure that the paths and conditions are specified correctly.

~~~
# ----------------------
# make metadata for BindingSiteFinder
# ----------------------
meta = data.frame(
  id = c(1,2), # give each sample a unique id
  condition = c(“cond_A”, “cond_A”), # add the condition for
each sample (used for differential analysis), but one BSF
object per condition needs to be used
 clPlus = bw.plus, # crosslinks plus strand bigwig files
 clMinus = bw.minus) # crosslinks minus strand bigwig files
meta
~~~
  e. Use the function BSFind() for automatic binding site generation. We recommend to turn off the gene-wise filter (cutoff.geneWiseFilter = 0) as automatic estimation for this parameter is overly stringent.

~~~
# ----------------------
# run BindingSiteFinder in automatic mode
# ----------------------
# make BSF object
bds_object = BSFDataSetFromBigWig(ranges = peaks,
                           meta = meta,
                           silent = TRUE)
# compute initial binding sites allowing BindingSiteFinder
to estimate most parameters
bds_automatic = BSFind(bds_object,
                           anno.genes = gns,
                           anno.transcriptRegionList = regions,
                           cutoff.geneWiseFilter = 0)
# save automatic binding sites
saveRDS(bds_automatic, paste0(out, “bds_automatic.rds”))
~~~
  f. Get a summary of BSF object including the number of binding sites (#N Ranges), the width of the binding sites (Width ranges) and the number of samples that were considered for binding site definition (#N Samples).

~~~
# binding site summary
bds_automatic
~~~
  g. Display the performed steps including the automatically estimated parameters in a workflow chart (**Fig. 7A**).

~~~
# visualize steps and filters
processingStepsFlowChart(bds_automatic)
~~~

**Figure 7.**
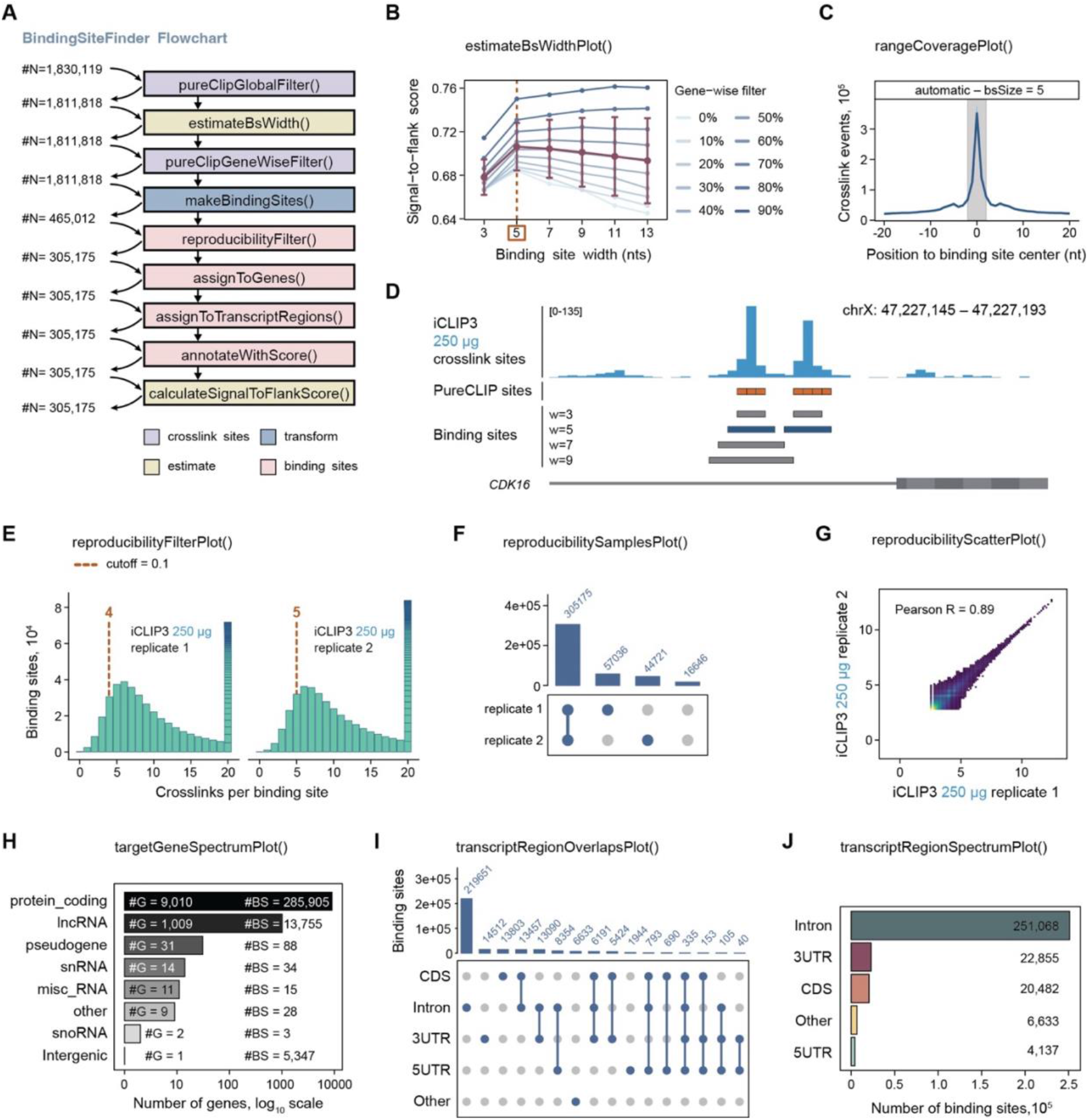
Definition of binding sites with BindingSiteFinder. Exemplary visualizations for the 250 µg U2AF2 iCLIP3 sample. **(A)** BindingSiteFinder workflow summary. Numbers indicate peak or binding site counts at each step before or after binding site definition, respectively. **(B–D)** Estimation of optimal binding site width. **(B)** estimateBsWidthPlot displays signal-to-flank scores (y-axis) across candidate widths (x-axis), calculated for ten peak subsets generated by gene-wise filtering (0–90%; light to dark blue). Mean and standard deviation per width are marked in dark red, and suggested optimal width (here: 5 nt) in orange dashed line. **(C)** rangeCoveragePlot depicts metaprofile of crosslink events (y-axis) in 20-nt window around binding site center (x-axis); the grey box indicates given width. **(D)** Genome browser view in the gene *CDK16* (chrX:47,227,145-47,227,193) with U2AF2 iCLIP3 crosslink signal (blue), PureCLIP peaks (orange) and binding sites of 3, 5, 7 and 9 nt width (grey /dark blue). **(E–G)** Reproducibility filtering. **(E)** reproducibilityFilterPlot shows number of binding sites (y-axis) with given number of crosslinks (x-axis) for each replicate with selected cutoff for a binding site being supported (repro.cutoff = 0.1; corresponding 4 and 5 crosslink events for replicates 1 and 2, respectively). **(F)** reproducibilitySamplesPlot displays intersections of supported sites between replicates. **(G)** reproducibilityScatterPlot shows crosslink counts per binding site (log2 scale) in replicate 1 (x-axis) versus replicate 2 (y-axis), colored by point density; Pearson correlation coefficient is indicated. **(H–J)** Assignment of binding sites to bound genes and transcript regions. **(H)** targetGeneSpectrumPlot summarizes bound genes by gene biotype, indicating gene biotype (left) and binding site counts (right). **(I)** transcriptRegionOverlapsPlot shows ambiguous assignments to multiple transcript regions. **(J)** Final distribution of binding sites across transcript regions after hierarchical assignment of ambiguous binding sites. An HTML report with the complete BindingSiteFinder analysis is provided in Supplementary Material 3.

#### Visual inspection of binding sites

7. Visually inspect the generated binding sites in a genome browser. **Note:** The following steps are given for the Integrative Genomics Viewer (IGV, https://igv.org/) but can be transferred to other genome browsers like the UCSC Genome Browser (https://genome.ucsc.edu/).
  a. Save the binding sites as BED file.

~~~
# export binding sites as BED file
exportToBED(bds_automatic, con = paste0(out,
“BindingSites_automatic.bed”))
~~~
  b. Load data in IGV.
    i. Open IGV and select your genome.
    ii. Load crosslink BIGWIG files from the folder bw_merged.
    iii. Turn of the windowing function (“None”) by right-click on the sample names.
    iv. Load peak BED file(s) with the PureCLIP-called peaks from the folder peaks.
    v. Load the BED file BindingSites_automatic.bed with the binding sites exported in the previous step.
  c. Navigate to genes of interest and/or housekeeping genes with binding sites and inspect the distribution of crosslink events within and around the binding sites. **Note:** With sufficient coverage, the crosslink events in a binding site usually follow a bell-shaped distribution. For comparison between binding sites (and between samples), keep in mind that the signal intensity both within the binding sites and in the surrounding background is proportional to the expression of the gene (and the sequencing depth of the sample).
    i. Binding site width: Do the defined binding sites accurately capture the spread of the crosslink events? Should the binding sites be wider or narrower? Refer to the section “Optimize binding site width” section to evaluate this further and possibly change the width accordingly.
    ii. Filter settings: Do many binding sites hardly rise over background or, inversely, are many prominent looking, bell-shaped patterns not called as binding sites? In this case, refer to the section “Selecting custom cutoffs”.

**Note:** Beware that not all binding sites will look perfect with any parameter settings. Try to find a combination of parameters for which the majority looks good.

#### Optimize binding site width

8. Evaluate how BindingSiteFinder decided for the optimal binding site width. In brief, BindingSiteFinder computes the signal-to-flank ratio, i.e., the ratio of crosslink events within the binding sites over equal-sized windows to both sides, for a range of widths (**Fig. 7B**). To account for different binding strengths, the ratios are averaged over calculations with increasing gene-wise filter, i.e., each time excluding an additional 10% of ranked PureCLIP-called peaks per gene (geneWiseFilter from 0 to 0.9). The optimal width is then chosen to maximize the median signal-to-flank ratio across all binding sites. **Note:** The width is always uneven because the binding sites are aligned to the highest signal in the center and then resized evenly to both sites (Busch et al., 2020).
  a. Use the function estimateBsWidthPlot() to check how BindingSiteFinder decided for the optimal binding site width.

~~~
# check estimation of binding site width
estimateBsWidthPlot(bds_automatic)
~~~
  b. Check whether other widths besides the chosen value reached almost equally high signal-to-flank ratios.
  c. Generate binding sites with alternative widths and compare them to the initially defined set (**Fig. 7C, D**).
    i. Call the function makeBindingSites() with a range of values for the parameter bsSize. The example code below implements binding sites of width 5, 7, and 9.
    ii. Use the function rangeCoveragePlot() to visualize the summed coverage of crosslink events across binding sites for each width (**Fig. 7C**).

~~~
# ----------------------
# compare different binding site widths
# ----------------------
# compute binding sites with width 5, 7, and 9
bds1 <- makeBindingSites(object = bds_object, bsSize = 5)
bds2 <- makeBindingSites(object = bds_object, bsSize = 7)
bds3 <- makeBindingSites(object = bds_object, bsSize = 9)
# summarize in list
l = list(‘automatic - bsSize = 5’ = bds1,
          ‘close by - bsSize = 7’ = bds2,
          ‘close by - bsSize = 9’ = bds3)
# plot comparison
rangeCoveragePlot(l, width = 20, show.samples = F,
subset.chromosome = “chr1”)
~~~
    iii. Evaluate how well the binding site boundaries (grey box) capture the crosslinking signal (blue line) which should display a peak in the middle and recede to background levels on both sides .
    **Note:** We chose a width of 5 for the U2AF2 data shown, but 7 would also be a valid option. If analyzing multiple CLIP data sets of the same protein, select the same width for all to allow for direct comparisons.
  d. Rerun BSFind() with the selected width (here: bsSize = 7).

~~~
# ----------------------
# optional: use a different width
# ----------------------
bds_selected_width = BSFind(bds_object,
                 anno.genes = gns,
                 anno.transcriptRegionList = regions,
                 bsSize = 7,
                 cutoff.geneWiseFilter = 0)
# check new number of binding sites
bds_selected_width
~~~
  e. Export the new binding sites as BED file and load them in IGV for visual inspection.

~~~
# save and export new binding sites
saveRDS(bds_selected_width, paste0(out, “
bds_selected_width.rds”))
exportToBED(bds_selected_width, con = paste0(out,
“BindingSites_resize.bed”))
~~~

#### Select custom cutoffs (optional)

BindingSiteFinder implements multiple additional parameters that can be used to fine-tune the criteria for binding site definition. For instance, the parameter geneWiseFilter allows to exclude a user-defined fraction of PureCLIP-called peaks per gene from the binding site definition steps. This can be useful for focusing on the most prominent binding sites per genes. For a full description of the available parameters, refer to the documentation of BindingSiteFinder at https://www.bioconductor.org/packages/release/bioc/html/BindingSiteFinder.html.

**Optional:** Set the gene-wise filter to 0.1 as an example.

~~~
# ----------------------
# optional: use a different gene-wise filter #
# ----------------------
bds_genewisefilter = BSFind(bds_object,
                 anno.genes = gns,
                 anno.transcriptRegionList = regions,
                 bsSize = 7,
                 cutoff.geneWiseFilter = 0.1)
# check new number of binding sites
bds_genewisefilter
~~~

#### Reproducibility filter

BindingSiteFinder uses a two-step procedure in which binding sites are first defined on the merged samples—thereby increasing the signal-to-background ratio—and then tested for reproducibility across the individual samples. Based on a user-defined percentile (repro.cutoff), the minimum number of crosslink events that are required to support a binding site is chosen individually for each sample based on its overall signal depth (Busch et al., 2020). Additionally, the minimum number of supporting samples can be set with the parameter repro.nReps. By default, BindingSiteFinder uses n-1 samples for repro.nReps, with repro.cutoff = 0.05 (see below).

**Note:** If only one sample is available (not recommended!), the default value n-1 for repro.nReps in BSFind() will not work. In this case, turn off reproducibility filtering by setting repro.nReps = 1 and repro.cutoff = 0.

9. Assess the reproducibility of binding sites across samples.
  a. Evaluate the signal distribution and minimum crosslink events per binding site chosen for each sample. The function reproducibilityFilterPlot() visualizes the distribution of crosslink events per binding site, indicating the number of crosslink events corresponding to the selected percentile cutoff (**Fig. 7E**).

~~~
# plot support cutoffs used for each sample
reproducibilityFilterPlot(bds_selected_width)
~~~ **Note:** The support cutoff is defined as a percentile, rather than an absolute number, to account for differences in signal depth.
  b. Evaluate how many binding sites are supported by a sufficient number of samples defined by repro.nReps (**Fig. 7F**). If only two replicates are available, we recommend requiring support from both (repro.nReps = 2). With more replicates, require support by all or all but one sample. n-1 samples or less is advisable if some samples are considerably worse to not lose too many binding sites.

~~~
# intersections of supported binding sites per sample
reproducibilitySamplesPlot(bds_selected_width)
~~~
  c. Re-run BSFind() with the selected reproducibility parameters. **Note:** A higher support cutoff of 0.1 or more can be useful if the data shows high background. However, in shallow data sets, higher reproducibility cutoffs can lead to an overly loss of binding sites.

~~~
# make binding sites using new reproducibility cutoffs
bds_repro = BSFind(bds_object,
                   anno.genes = gns,
                   anno.transcriptRegionList = regions,
                   bsSize = 5, # add the width you selected
                   cutoff.geneWiseFilter = 0,
                   repro.nReps = 2,
                   repro.cutoff = 0.1)
# get numbers
bds_repro
# plot reproducibility support with changed settings
reproducibilityFilterPlot(bds_repro)
~~~
  d. Save the final binding sites as RDS file and export them as a BED file.

~~~
# get final binding sites
bds_final <- bds_selected_width
# save and export
saveRDS(bds_final, paste0(out, “bds_final.rds”))
exportToBED(bds_final, con = paste0(out, “binding_sites_final.bed”))
~~~
  e. Load the binding sites as BED file in IGV. To evaluate reproducibility, load the crosslinks (BIGWIG files) of the individual samples for comparison.

#### Characterization of binding sites

10. Assign binding sites to target genes. **Note:** In regions of overlapping annotations, binding sites cannot be unambiguously assigned to a single gene. One possibility to resolve such overlaps in BindingSiteFinder is by applying a hierarchy of gene biotypes. Since CLIP data is strand-specific; this problem does not apply for genes on opposite strands.
  a. Inspect the gene biotypes that are present in the annotation.

~~~
# present gene types
unique(gns$gene_type)
~~~ The GENCODE annotation used here contains the following biotypes: protein_coding transcribed_unprocessed_pseudogene, processed_pseudogene, lncRNA, transcribed_unitary_pseudogene, transcribed_processed_pseudogene, unprocessed_pseudogene, IG_V_pseudogene, unitary_pseudogene, TR_V_pseudogene, IG_V_gene, snRNA, miRNA, misc_RNA, snoRNA, rRNA_pseudogene, rRNA, vault_RNA, TR_V_gene, Mt_tRNA, Mt_rRNA, IG_C_gene, IG_J_gene, TR_J_gene, TR_C_gene, TR_J_pseudogene, IG_D_gene, ribozyme, IG_C_pseudogene, TR_D_gene, TEC IG_J_pseudogene, scaRNA, translated_processed_pseudogene, artifact, sRNA, IG_pseudogene.
  b. Visualize the number of binding sites that overlap with multiple gene biotypes.

~~~
# binding sites with overlapping gene biotypes
geneOverlapsPlot(bds_final)
~~~
  c. Select which gene biotypes should be kept separately, depending on your protein of interest. Then, collapse all pseudogene categories into “pseudogenes” and all remaining categories into “other”.

~~~
# decide which gene biotypes to keep separately
my_gene_types <- c(“protein_coding”, “lncRNA”, “snRNA”,
“snoRNA”, “miRNA”, “rRNA”, “misc_RNA”, “tRNA”)
~~~

~~~
# make hierarchy column
gns <- as.data.frame(gns) %>%
   dplyr::mutate(gene_type_plot = case_when(
      gene_type %in% my_gene_types ∼ gene_type,
      grepl(gene_type, pattern = “pseudogene”) ∼ “pseudogene”,
      TRUE ∼ “other”
   )) %>%
   makeGRangesFromDataFrame(., keep.extra.columns = TRUE)
~~~
  d. Define the hierarchy of the gene biotypes with a vector sorted by decreasing relevance.

~~~
# define hierarchy of interesting gene biotypes
hierarchy <- c(“protein_coding”, “lncRNA”, “snRNA”,
“snoRNA”, “miRNA”, “rRNA”, “misc_RNA”, “tRNA”, “pseudogene”,
“other”)
~~~
  e. Apply the hierarchy and plot the resulting distribution of binding sites across the gene biotypes (**Fig. 7H**).

~~~
# assign gene type according to hierarchy
bds_alt_gene_asignment <-
assignToGenes(bds_final,
                        overlaps = “hierarchy”,
                        overlaps.rule = hierarchy,
                        anno.genes = gns,
                        match.geneType = “gene_type_plot”
                        )
# plot
targetGeneSpectrumPlot(bds_alt_gene_asignment, showNGroups =
20)
~~~
11. Assign the binding sites to the transcript regions in which they are located. As most genes contain multiple transcript isoforms, overlaps are again resolved with a hierarchy.
  a. Visualize the amount of ambiguous binding sites overlapping with multiple transcript regions (**Fig. 7I**).

~~~
# visualize binding sites overlapping with multiple
transcript regions
transcriptRegionOverlapsPlot(bds_final)
~~~
  b. Select a hierarchy which best fits to the expected binding behavior of your protein of interest. For U2AF2 as a known splicing factor, we give the highest priority to introns, followed by the untranslated regions and the coding region.

~~~
# transcript region hierarchy
region_hierarchy <- c(“INTRON”, “UTR5”, “UTR3”, “CDS”)
~~~ **Note:** The names of the regions need to be identical to the names assigned when preparing the transcript regions object from the annotation.
  c. Assign the binding sites to the transcript regions and plot the resulting distribution (**Fig. 7J**).

~~~
# assign binding sites to transcript regions by the chosen
hierarchy
bds_final <-
assignToTranscriptRegions(bds_final,
                          overlaps = “hierarchy”,
                          overlaps.rule = region_hierarcy,
anno.transcriptRegionList = regions)
# plot
transcriptRegionSpectrumPlot(bds_final, show.others = TRUE)
~~~

### EXPECTED OUTCOMES

As a showcase example, we defined U2AF binding sites from the iCLIP3 data from HeLa cell lysates containing 250 µg, 100 µg, and 40 µg of total protein (see above). Selected plots from the BindingSiteFinder analysis for the 250 µg U2AF2 samples are shown in **Fig. 7**, including the workflow summary, the estimation of the optimal binding site width, the assessment of reproducibility and the assignment of binding sites to bound genes and transcript regions. An HTML report showing the complete BindingSiteFinder analysis including diagnostic plots and other visualizations is provided in **Supplementary Material 3**.

Comparison between the 250 µg, 100 µg, and 40 µg iCLIP3 samples shows that— irrespective of the amount of input material—a similar library depth was reached (**Fig. 8A**). The binding patterns show a strong qualitative and quantitative agreement across the iCLIP3 samples and to a previously published U2AF2 iCLIP2 dataset in the same cell line (**Fig. 8B–E**).

**Figure 8.**
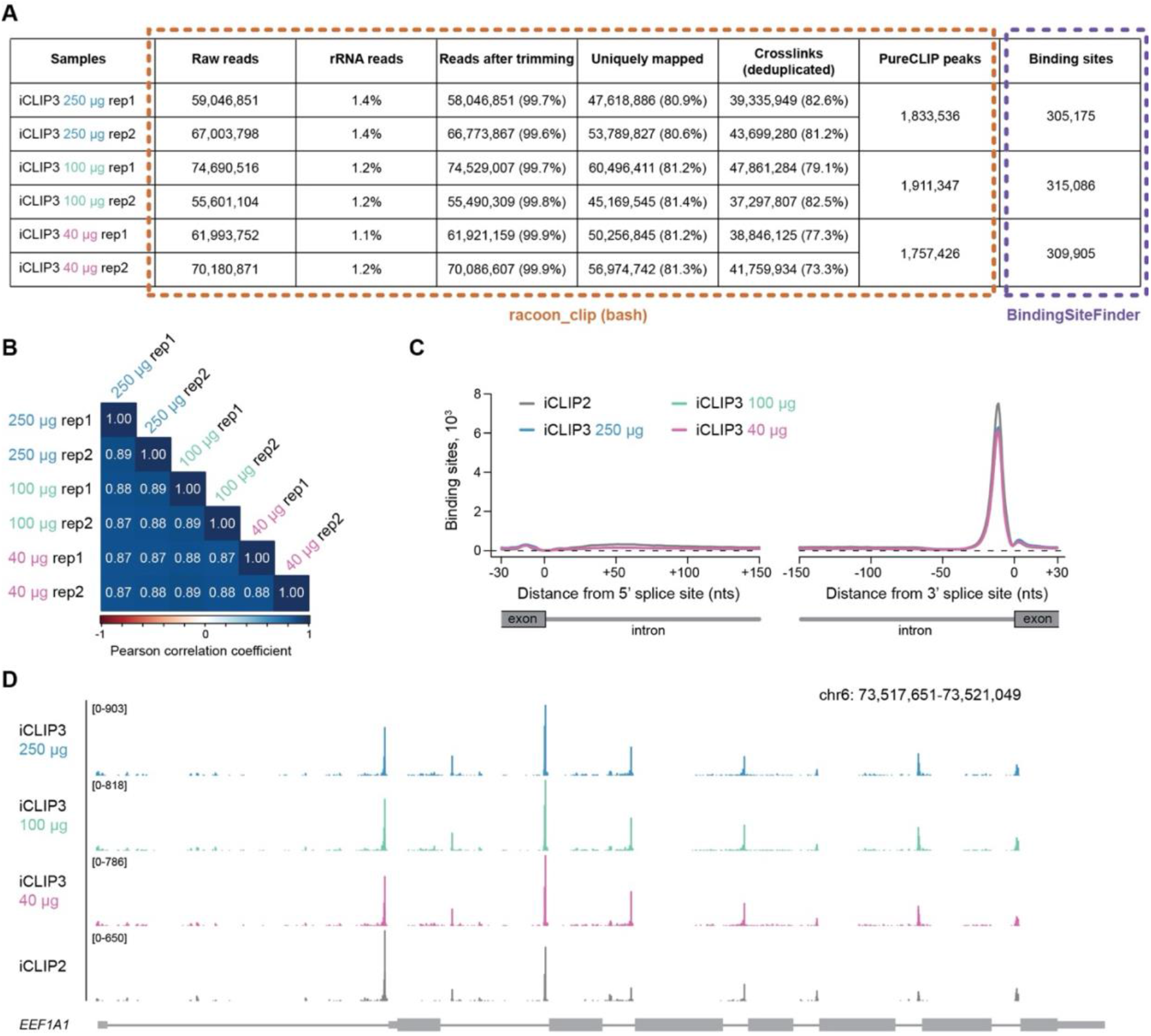
Comparison between U2AF2 iCLIP3 samples. Data were individually processed for U2AF2 iCLIP3 from HeLa cell lysates containing 250 µg, 100 µg, and 40 µg of total protein. **(A)** Number of sequenced and mapped reads, PureCLIP-called peaks (from racoon_clip) and binding sites (from BindingSiteFinder) for all U2AF2 iCLIP3 samples. Percentages correspond to the difference from the prior step. Peak calling and binding site definition were performed on the merge of replicates for each condition. **(B)** Pairwise Pearson correlation coefficients for all replicates based on the number of crosslink events in overlapping binding sites. **(C)** Metaprofile of binding sites around 5′ and 3′ splice sites (smoothed lines). blue – iCLIP3 250 µg, green - iCLIP3 100 µg, pink – iCLIP3 40 µg, grey – published U2AF2 iCLIP2 data (GSM6793346, GSM6793347) (Ebersberger et al., 2023). **(D)** Genome browser view of U2AF2 crosslink events (sum of replicates) across the gene *EEF1A1* (chr6:73,517,651-73,521,049) of U2AF2 iCLIP3 and iCLIP2 experiments. Axis scale is depicted on the left.

For the U2AF2 iCLIP3 data in HeLa cells, we detected more than 35 million crosslink events per sample. Although this number cannot be generalized, at least 1 million crosslink events can be expected—as a rule of thumb—for good RNA-binders in mammalian cells. If the signal depth of an iCLIP3 dataset is much lower, we recommend revisiting the quality controls performed during the experiment and data analysis. Similarly, detecting less than 1.000 binding sites may point to limited signal depth in the data.

### LIMITATIONS

Defining a hierarchy of gene biotypes and transcript regions requires prior knowledge about the protein of interest and can potentially skew the resulting distribution. To avoid such biases, orient the hierarchy on distribution of uniquely assigned binding sites or use a majority vote (see BindingSiteFinder documentation for more details). Moreover, for the interpretation of bound transcript regions, it is important to keep in mind that some regions, particularly introns, are considerably longer than others.

Finally, overlap between biological replicates or independent experiments is never expected to be complete. Variability in crosslink efficiency, sequencing depth, library complexity, and peak calling contributes to incomplete overlap even under well-controlled conditions. Therefore, reproducibility should be assessed quantitatively and interpreted in the context of experimental noise rather than assuming perfect overlap of binding sites.

### TROUBLESHOOTING

#### Problem 1

A high number of binding sites is removed in the reproducibility filter step.

#### Potential solution

Check in the reproducibilityFilterPlot whether certain samples show much lower signal than others. To avoid that weak samples impair the overall reproducibility, reduce the minimum number of samples required to support a binding site or completely remove the given samples from the analysis.

#### Problem 2

The binding site definition with BindingSiteFinder fails.

#### Potential solution

A possible reason is that the number of crosslink events and/or PureCLIP-called peaks is too low. This can sometimes be circumvented by turning off the automatic estimation of binding width by directly setting the parameter bsSize to the desired value. Go back to the racoon_clip report to evaluate data quality across the analysis.

#### Problem 3

Presumed nice peaks are not captured as binding sites.

#### Potential solution

This may result from a lack of reproducibility (see above) or a generally low signal in the region, usually occurring on lowly expressed transcripts. As a rule of thumb, peaks below 20 crosslink events are close to background levels and not always found by PureCLIP.

### KEY RESOURCES TABLE

#### Reagents

| REAGENT or RESOURCE | SOURCE | IDENTIFIER |
| --- | --- | --- |
| Nuclease-free water | ThermoFisher Scientific | 10977035 |
| 1 M Tris-HCl, pH 7.5 | ThermoFisher Scientific | 15567027 |
| 5 M Sodium Chloride (NaCl) | ThermoFisher Scientific | AM9759 |
| Sodium deoxycholate, powder | Merck/Sigma | D6750-100G |
| Igepal CA-630 | Merck/Sigma | I8896-50ml |
| Tween-20 | Merck/Sigma | P9416-50ml |
| 10% SDS solution | ThermoFisher Scientific | 24730020 |
| 1 M Magnesium Chloride (MgCl<sub>2</sub>) | ThermoFisher Scientific | AM9530G |
| 0.5 M EDTA, pH 8.0 | ThermoFisher Scientific | AM9260G |
| 1 M Tris-HCl, pH 6.5 | Merck/Sigma | 20-160 |
| 1 M DL-Dithiothreitol (DTT in H<sub>2</sub>O) | Merck/Sigma | 646563-10X.5ml |
| Dynabeads Protein G for IP | ThermoFisher Scientific | 10003D |
| cOmplete Mini, EDTA-free Protease Inhibitor Cocktail | Merck/Sigma | 11836170001 |
| Ambion RNase I, cloned (100 U/μl) | ThermoFisher Scientific | AM2294 |
| TURBO DNase (2 U/μl) | ThermoFisher Scientific | AM2238 |
| RNaseIn Ribonuclease Inhibitor | Promega | N2511 |
| T4 Polynucleotide kinase (PNK) | New England Biolabs | M0201S |
| T4 RNA Ligase 1, high concentration | New England Biolabs | M0437M |
| 10X T4 RNA Ligase Reaction buffer | New England Biolabs | in M0437M |
| 50% Polyethylene Glycol (PEG) 8000 | New England Biolabs | in M0437M |
| 100 mM ATP | New England Biolabs | in M0437M |
| 1 mM pCp-IR750 | Jena Bioscience | NU-1706-IR750 |
| 4-12% NuPAGE Bis-Tris Protein gels | ThermoFisher Scientific | NP0322BOX |
| 20X NuPAGE MOPS SDS Running buffer | ThermoFisher Scientific | NP0001 |
| 20X NuPAGE Transfer buffer | ThermoFisher Scientific | NP00061 |
| 4X NuPAGE LDS Sample buffer | ThermoFisher Scientific | NP0007 |
| PageRuler Prestained protein ladder | ThermoFisher Scientific | 26616 |
| Proteinase K | Merck/Sigma | 3115828001 |
| Phenylmethylsulfonylfluoride (PMSF) | Cell Signaling | 8553S |
| RNA Clean and Concentrator 5 kit | Zymo Research | R1015 |
| Methanol | Carl Roth | 0082.1 |
| Ethanol | Merck/Sigma | 32205-1L-M |
| Isopropanol | Merck/Sigma | 190764 |
| SuperScript III Reverse Transcriptase | ThermoFisher Scientific | 18080044 |
| 5X First Strand buffer | ThermoFisher Scientific | in 18080044 |
| 0.1 M DTT | ThermoFisher Scientific | in 18080044 |
| dNTP Solution mix (10 mM each) | New England Biolabs | N0447S |
| 1 M HEPES solution | Merck/Sigma | H0887-20 ml |
| Sodium Hydroxide (NaOH) | Merck/Sigma | S8045-500G |
| Dynabeads MyOne Silane | ThermoFisher Scientific | 37002D |
| Buffer RLT | Qiagen | 79216 |
| 2X Phusion High-Fidelity PCR MasterMix | New England Biolabs | M0531S |
| DMSO | New England Biolabs | in M0531S |
| GeneRuler Ultra Low Range DNA ladder | ThermoFisher Scientific | SM1211 |
| ProNex Size Selective Purification System | Promega | NG2001 |
| 2X TBE-Urea Sample buffer | ThermoFisher Scientific | LC6876 |
| Low Range ssRNA ladder | New England Biolabs | N0364S |
| 10X TBE buffer, RNase-free (1L) | ThermoFisher Scientific | AM9863 |
| 10% Novex TBE-Urea gels | ThermoFisher Scientific | EC68752BOX |
| SYBR Gold (10,000X) | ThermoFisher Scientific | S11494 |

#### Equipment

| REAGENT or RESOURCE | SOURCE | IDENTIFIER |
| --- | --- | --- |
| UV crosslinker | Vilber | 24-2460 |
| DynaMag-2 magnetic rack (1.5 ml tubes) | ThermoFisher Scientific | 12321D |
| Magnetic separation rack (0.2 mL tubes) | BioCat | 10-0008-EPC |
| Proteus clarification mini spin columns | SERVA Electrophoresis | 42225.01 |
| XCell SureLock and XCell II modules | ThermoFisher Scientific | EI0002 |
| Whatman filter paper | GE Healthcare | 3030917 |
| Protran nitrocellulose membrane BA85 | Cytiva | 10600002 |
| Swann-Morton surgical scalpels | Fisher Scientific | 11728353 |
| PowerPac Basic Power Supply | Bio-Rad | 1645050 |
| ChemiDoc MP Imaging System | Bio-Rad | N/A |
| Thermomixer | N/A | N/A |
| PCR tubes | Sarstedt | 72.991.002 |
| 1.5 mL DNA Low binding tubes | Sarstedt | 72.706.700 |
| Non-stick RNase-free 1.5 mL tubes | ThermoFisher Scientific | AM12450 |
| PCR Thermocycler | N/A | N/A |
| Qubit Fluorimeter | ThermoFisher Scientific | Q33238 |
| Tape Station 4150 or 4200 | Agilent Technologies | N/A |
| High Sensitivity D1000 Screen Tape | Agilent Technologies | 5067-5584 |
| High Sensitivity D1000 Reagents | Agilent Technologies | 5067-5585 |
| Tape Station tube strips | Agilent Technologies | 401428 |
| Tape Station tube caps | Agilent Technologies | 401425 |

#### Software and algorithms

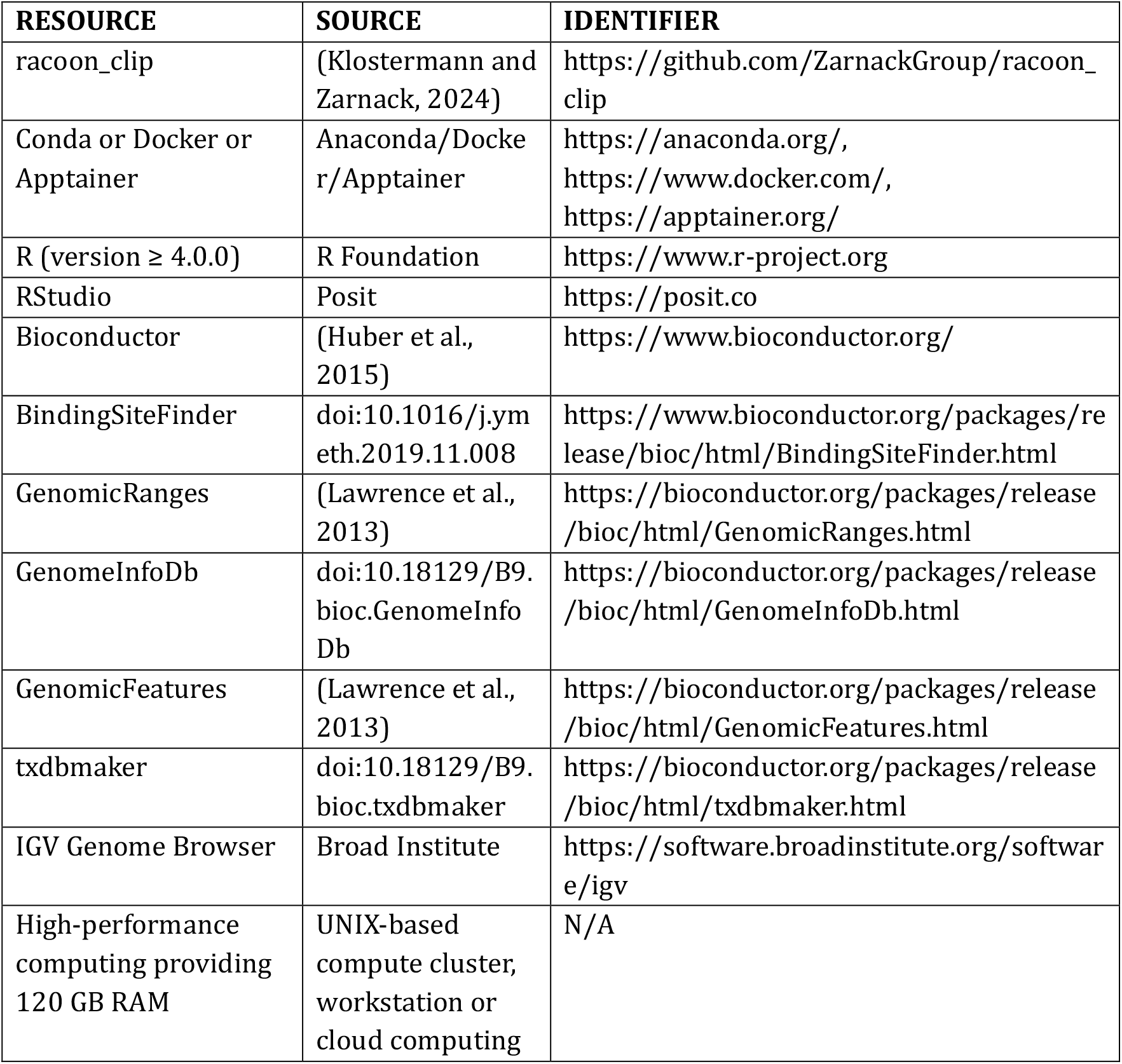

| RESOURCE | SOURCE | IDENTIFIER |
| --- | --- | --- |
| racoon_clip | (Klostermann and Zarnack, 2024) | https://github.com/ZarnackGroup/racoon_clip |
| Conda or Docker or Apptainer | Anaconda/Docker/Apptainer | https://anaconda.org/,<br>https://www.docker.com/,<br>https://apptainer.org/ |
| R (version $\geq 4.0.0$) | R Foundation | https://www.r-project.org |
| RStudio | Posit | https://posit.co |
| Bioconductor | (Huber et al., 2015) | https://www.bioconductor.org/ |
| BindingSiteFinder | doi:10.1016/j.ymeth.2019.11.008 | https://www.bioconductor.org/packages/release/bioc/html/BindingSiteFinder.html |
| GenomicRanges | (Lawrence et al., 2013) | https://bioconductor.org/packages/release/bioc/html/GenomicRanges.html |
| GenomeInfoDb | doi:10.18129/B9.bioc.GenomeInfoDb | https://bioconductor.org/packages/release/bioc/html/GenomeInfoDb.html |
| GenomicFeatures | (Lawrence et al., 2013) | https://bioconductor.org/packages/release/bioc/html/GenomicFeatures.html |
| txdbmaker | doi:10.18129/B9.bioc.txdbmaker | https://bioconductor.org/packages/release/bioc/html/txdbmaker.html |
| IGV Genome Browser | Broad Institute | https://software.broadinstitute.org/software/igv |
| High-performance computing providing 120 GB RAM | UNIX-based compute cluster, workstation or cloud computing | N/A |

#### Deposited data

| RESOURCE | SOURCE | IDENTIFIER |
| --- | --- | --- |
| Genome assembly (e.g., GRCm38 / hg38) | GENCODE | https://www.encodegenes.org |
| Gene annotation (e.g., GENCODE, here release 49) | GENCODE | https://www.encodegenes.org |
| (optional) rRNA sequences | NCBI/Refseq | https://www.ncbi.nlm.nih.gov/nucleotide/ |
| (optional) Genomes of potential contaminating organisms | ENSEMBL | https://www.ensembl.org/index.html |

#### Oligonucleotides

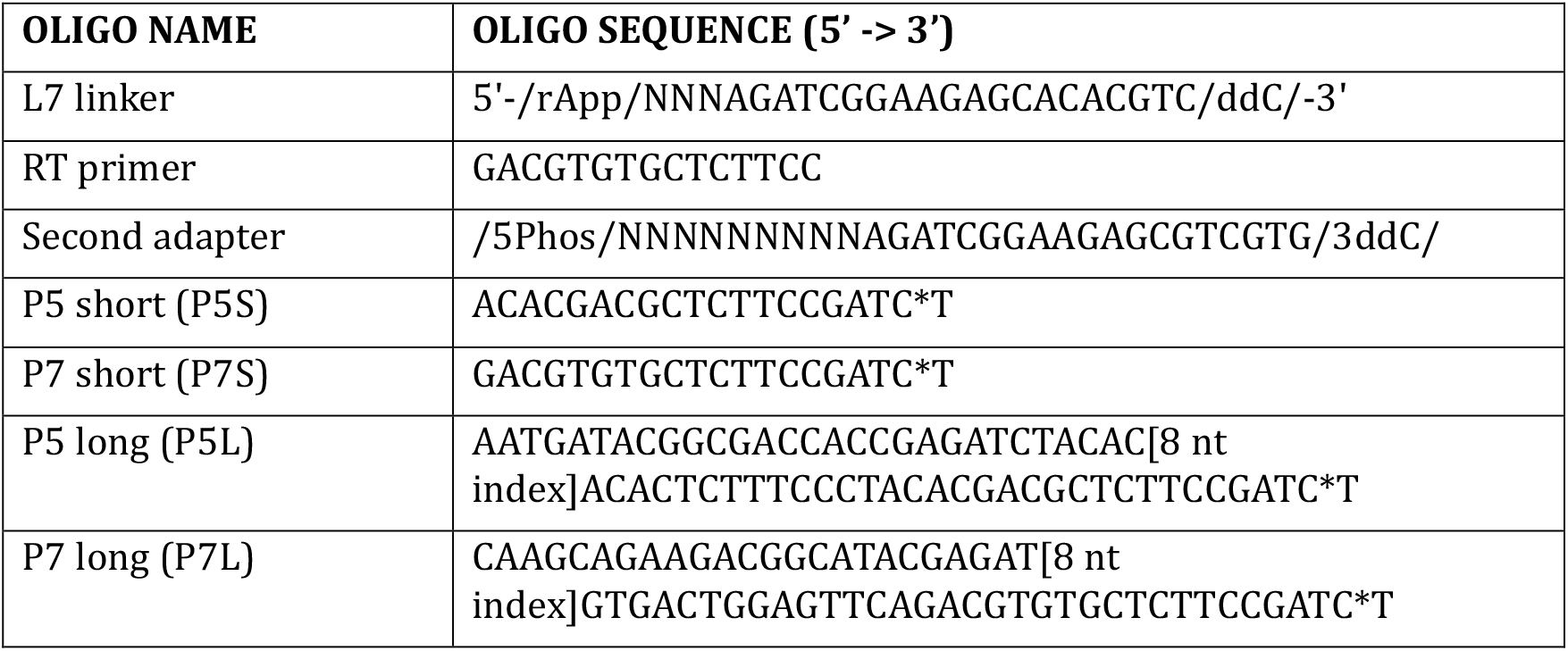

#### Reagent setup

**Lysis buffer**

50 mM Tris-HCl pH 7.5, 100 mM NaCl, 1% (vol/vol) Igepal CA-630, 0.5% (wt/vol) Sodium deoxycholate, 0.1% SDS (vol/vol)

**High Salt buffer**

50 mM Tris-HCl pH 7.5, 1 M NaCl, 1% (vol/vol) Igepal CA-630, 0.5% (wt/vol) Sodium deoxycholate, 0.1% SDS (vol/vol), 1 mM EDTA

**PNK Wash buffer**

20 mM Tris-HCl pH 7.5, 10 mM MgCl_2_, 0.2% (vol/vol) Tween-20

**Last Wash buffer**

20 mM Tris-HCl pH 7.5, 100 mM NaCl, 0.2% (vol/vol) Tween-20, 2 mM EDTA

**5X PNK buffer pH 6.5**

350 mM Tris-HCl pH 6.5, 50 mM MgCl_2_, 5 mM DTT

Freeze aliquots of the buffer. Do not thaw and freeze again.

**Proteinase K buffer**

100 mM Tris-HCl pH 7.5, 50 mM NaCl, 10 mM EDTA, 0.2% SDS (vol/vol)

**PEG8000/NaCl Mix**

20% PEG8000 (vol/vol), 2.5 M NaCl

## RESOURCE AVAILABILITY

### Materials availability

This study did not produce new materials, reagents, or cell lines.

### Data and code availability

The U2AF2 iCLIP3 data generated in this study have been deposited at NCBI Gene Expression Omnibus (GEO) and will be publicly available as of the date of publication. The U2AF2 iCLIP2 data used for comparison are available under the accession numbers GSM6793346 and GSM6793347.

Scripts and configuration files used for processing the U2AF2 iCLIP3 data in this publication are available on GitHub (https://github.com/ZarnackGroup/Despic_et_al_2026).

## ACKNOWLEDGMENTS

This research was supported by the German Research Foundation (Deutsche Forschungsgemeinschaft, DFG) through grants ZA 881/6-1 (to K.Z.), TRR 267 (project number 403584255, TP A03, to M.M.-M. and K.Z.), SFB 1551 (project number 464588647) to J.K.), the excellence cluster Cardiopulmonary Institute (CPI) EXS2026 (project number 390649896, to M.M.-M. and K.Z.), the excellence cluster SCALE EXS3094 (project number 533751785, to M.M.-M.), and the Cluster for Nucleic Acid Sciences and Technologies – NUCLEATE EXC 3113/1 (project number 533767322, to J.K. and K.Z.), as well as by EMBO (pc25/22, to Z.K., J.K. and M.M.-M.).

We gratefully acknowledge the Genomics Core Facility at Institute of Molecular Biology (IMB) Mainz for sequencing the U2AF2 iCLIP3 libraries. We would like to thank Miona C orovic, Elias Bechara, Rosario Avolio and all EMBO iCLIP course participants for successfully testing the experimental iCLIP3 protocol, Annika Ladwig and Sarah Wolf for testing the data analysis protocol, and all members of the Mu ller-McNicoll, Ko nig and Zarnack labs for discussion.

## AUTHOR CONTRIBUTIONS

V.D., J.K. and M.M.-M. conceived the project. V.D. developed the experimental protocol, with contributions from A.O. and M.M.. V.D. performed the experiments. M.K. performed the bioinformatics data analysis. V.D. and M.K. prepared the figures. A.B. developed the initial pipeline for processing iCLIP3 sequencing reads. A.O. performed the U2AF2 iCLIP3 experiment in the Ko nig lab. M.M. developed the Shiny app for library multiplexing with long primers with UDIs. V.D. wrote the experimental protocols and M.K. wrote the bioinformatics protocols, with input from all co-authors. K.Z., J.K. and M.M.-M. supervised the project.

